# Syncrip/hnRNPQ is required for activity-induced Msp300/Nesprin-1 expression and new synapse formation

**DOI:** 10.1101/585679

**Authors:** Josh Titlow, Francesca Robertson, Aino Järvelin, David Ish-Horowicz, Carlas Smith, Enrico Gratton, Ilan Davis

## Abstract

Memory and learning involve activity-driven expression of proteins and cytoskeletal reorganisation at new synapses, often requiring post-transcriptional regulation a long distance from corresponding nuclei. A key factor expressed early in synapse formation is Msp300/Nesprin-1, which organises actin filaments around the new synapse. How Msp300 expression is regulated during synaptic plasticity is not yet known. Here, we show that the local translation of *msp300* is promoted during activity-dependent plasticity by the conserved RNA binding protein Syncrip/hnRNP Q, which binds to *msp300* transcripts and is essential for plasticity. Single molecule imaging shows that Syncrip is associated *in vivo* with *msp300* mRNA in ribosome-rich particles. Elevated neural activity alters the dynamics of Syncrip RNP granules at the synapse, suggesting a change in particle composition or binding that facilitates translation. These results introduce Syncrip as an important early-acting activity-dependent translational regulator of a plasticity gene that is strongly associated with human ataxias.

**Syncrip regulates synaptic plasticity via *msp300*:** Titlow et al. find that Syncrip (hnRNPQ RNA binding protein) acts directly on *msp300* to modulate activity-dependent synaptic plasticity. *In vivo* biophysical experiments reveal activity-dependent changes in RNP complex sizes compatible with an increase in translation at the synapse.

## Introduction

Activity-dependent neuronal plasticity is the cellular basis of memory and learning, involving the formation of new synapses and cytoskeletal remodeling in response to neuronal activity (West and Greenberg, 2011). To achieve plasticity, it is thought that neuronal activation leads to the elevated expression of over 1,000 different genes. Many activity-dependent genes have been identified either through RNA sequencing studies (Chen et al., 2016) or proteomics analysis (Dieterich and Kreutz, 2016) and the majority of the effort in the field has focused on explaining how activity leads to changes in gene expression through transcriptional regulation (Madabhushi and Kim, 2018). However, activity-dependent plasticity often occurs too rapidly and too far away from the cell nucleus to be explained by *de novo* transcription alone. Therefore, post-transcriptional regulation is thought to be a crucial mechanism to explain changes in gene expression in response to neuronal activity.

During synaptic plasticity, the actin cytoskeleton is extensively remodeled, a process requiring numerous regulatory proteins (Spence and Soderling, 2015). Nesprins are an especially interesting class of actin regulatory proteins because they connect both synapses and nuclei through the cytoskeleton. The Nesprins are encoded by *synaptic nuclear envelope-1* and *−2* (*SYNE-1* and *−2*), which contain at least 80 disease-related variants that cause cerebellar ataxias or muscular dystrophies (Zhou et al., 2018b). The molecular function of Nesprins and their role in muscular diseases are relatively well studied in mouse models of *SYNE-1* and *SYNE-2* (Zhou et al., 2018a) in relation to nucleo-cytoplasmic and cytoskeletal organization and function, but the function of Nesprins in neurological disorders is not yet known.

The synaptic function of Nesprins has begun to be investigated in the *Drosophila* orthologue, Msp300, one of many molecular components that are conserved between the *Drosophila* larval neuromuscular junction (NMJ) and mammalian glutamatergic synapses (Harris and Littleton, 2015; Menon et al., 2013; Titlow and Cooper, 2018) Msp300 is required for activity-dependent plasticity at the larval NMJ (Packard et al., 2015), where it organizes a post-synaptic actin scaffold around newly formed synapse clusters, known as boutons. The post-synaptic actin scaffold regulates glutamate receptor density at the synapse (Blunk et al., 2014), which also requires Msp300 (Morel et al., 2014). Msp300 is barely detectable at mature NMJ synapses but becomes highly enriched at the post-synapse in response to elevated neural activity (Packard et al., 2015). However, the mechanism by which activity-dependent Msp300 expression is regulated is poorly understood.

We previously identified *msp300* by RNA-Immunoprecipitation-Sequencing (RIP-Seq) to have the strongest interaction with an RNA binding protein (RBP) called Syncrip (Syp) (McDermott et al., 2014). The mammalian orthologue of Syp is hnRNP Q, an RBP that functions in a number of diverse biological processes ranging from sorting miRNA in exosome vesicles (Santangelo et al., 2016) to controlling the myeloid leukemia stem cell program (Vu et al., 2017), and was recently identified in a patient whole-exome sequencing study as a potential gene candidate for intellectual disability (Lelieveld et al., 2016). Syp is expressed throughout the mammalian brain (Tratnjek et al., 2017) and has been found in ribonucleoprotein (RNP) particles with FMRP protein (Chen et al., 2012), IP3 mRNA (Bannai et al., 2004) and BC200 mRNA (Duning et al., 2008). Knockdown of Syp in rat cortical neurons throughout development increases neurite complexity and alters the localisation of proteins encoded by its mRNA targets (Chen et al., 2012). Syp has also been shown to regulate the stability of its mRNA targets in macrophages (Kuchler et al., 2014). In the *Drosophila* larval neuromuscular junction (NMJ) Syp is expressed post-synaptically where it acts as a negative regulator of synapse development (Halstead et al., 2014; McDermott et al., 2014). Syp is required to maintain the correct synaptic pool of glutamatergic vesicles and therefore glutamatergic transmission at the larval NMJ. However, it is not known whether Syp regulates synapse formation or Msp300 expression in the context of activity-dependent synaptic plasticity.

Here, we show that Syp is required directly for synaptic plasticity and for regulating activity-induced Msp300 expression at the larval NMJ. Syp and *msp300* mRNA physically interact *in vivo* near the synapse in ribosome-containing granules, which become significantly less dynamic in response to elevated synaptic activity. Our work reveals a new RBP regulator that links neuronal activity to post-transcriptional regulation of an actin-regulating gene that is required for new synapse formation.

## Results

### Baseline and activity-dependent expression of Msp300 are post-transcriptionally regulated by Syp

In response to neuronal activity, Msp300 is rapidly enriched at the larval neuromuscular junction where it is required for structural synaptic plasticity (Fig. S1; (Packard et al., 2015)). We have also shown that *msp300* mRNA is associated with Syp protein in immunoprecipitation experiments using whole larval lysates (McDermott et al., 2014). To determine whether Msp300 expression is regulated by Syp at the larval NMJ, we quantified *msp300* mRNA and protein levels in *syp* mutant fillet preparations and compared them to wild type controls. We find that Msp300 expression at the larval NMJ is regulated post-transcriptionally by Syp. Primary *msp300* transcripts and cytosolic mRNA molecules were quantified using single molecule fluorescence *in situ* hybridization (smFISH; see Fig. S2 for details on probe design and controls). To test for an effect of Syp on *msp300* levels we used a previously characterized *syp* null allele, *syp*^*e00286*^ (McDermott et al., 2014). The spatial distribution of *msp300* transcripts in *syp*^*e00286*^ muscles was indistinguishable from wild type muscles (Fig. 1A). The number of primary transcripts (relative intensity of transcription foci) and the number of mature transcripts were also not significantly altered in *syp*^*e00286*^ relative to wild type larvae (Fig. 1B). Therefore, loss of *Syp* does not affect mRNA localization or mRNA turnover. To determine whether Msp-300 protein levels were affected by loss of Syp, we quantified Msp300 protein expression in larval muscles using immunofluorescence. Msp300 protein levels were on average 40% lower in *syp*^*e00286*^ than in wild type controls (Fig. 1C,D), which is in agreement with western blot data from whole larvae showing that Syp is a positive regulator of Msp300 translation during NMJ development (McDermott et al., 2014). Taken together with our smFISH data, these results suggest that proper translation of *msp300* transcripts in larval muscle requires Syp.

**Fig. 1.**
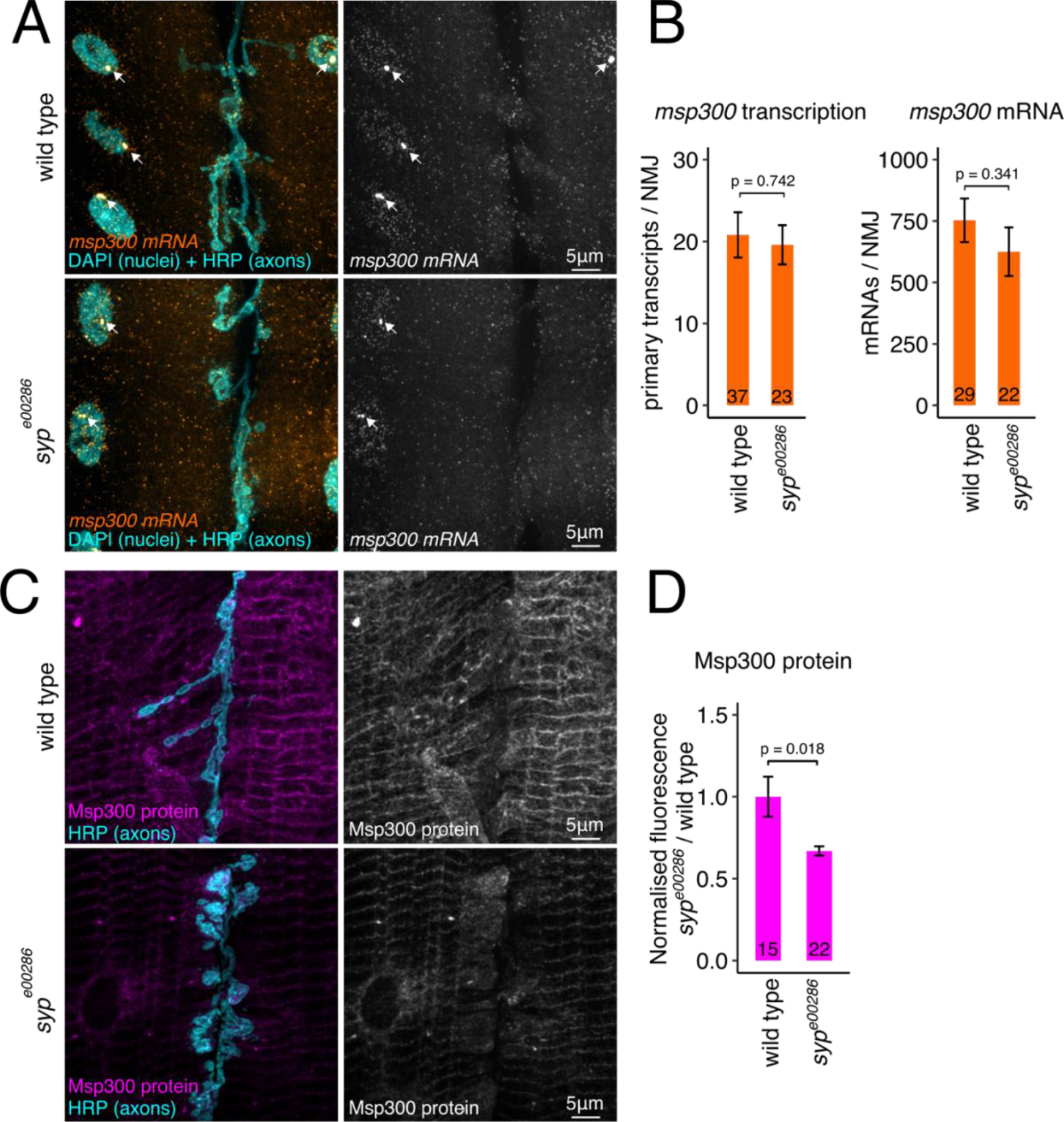
Msp300 expression at the larval NMJ is regulated post-transcriptionally by Syp. (A) *msp300* transcription and mRNA turnover are unaffected by loss of Syp. Single molecule fluorescence *in situ* hybridisation (smFISH) images show that steady state levels of *msp300* transcription (arrows) and cytosolic mRNA at wild type and *syp*^*e00286*^ mutant NMJs are similar. Quantification shows that loss of Syp does not have a significant effect on the level of primary or mature *msp300* transcripts at the larval NMJ. (C) Syp modulates Msp300 protein levels in larval muscle. Max z-projections of immunofluorescence images show that Msp300 protein levels in the muscles of *syp*^*e00286*^ mutant are reduced relative to wild type larvae. (D) Quantification of Msp300 immunofluorescence shows that Msp300 protein levels are significantly reduced in *syp*^*e00286*^ relative to wild type. Mean ± sem; Student’s unpaired T-test; number of NMJs measured shown in each bar.

To determine how activity-dependent enrichment of Msp300 at the synapse is regulated we measured the effect of Syp on Msp300 protein and mRNA levels in KCl stimulated samples, a well characterized model for synaptic plasticity (Fig. 2A). We found that Syp is required for post-synaptic enrichment of Msp300 in response to elevated synaptic activity. Msp300 protein levels increased by ~40% in stimulated NMJs relative to mock-treated controls (Fig. 2B,C). To determine if Syp is involved in elevating Msp300 levels, the experiment was repeated in the context of conditional Syp knockdown using the tripartite Gal80^ts^/Gal4/UAS system (Suster et al., 2004; see Methods for experimental details and Fig. S3 for controls). Conditional knockdown allowed us to separate potential developmental effects of Syp from specific activity-dependent effects. With Syp knocked down, the activity-dependent increase of Msp300 protein levels in the muscle was almost completely abolished (Fig. 2D,E). *msp300* mRNA levels were not significantly affected by neural activity (Fig. 2F,G), indicating that Syp does not act through mRNA turnover. We conclude that Syp is required directly for elevating Msp300 protein levels in response to increased neural activity level at the NMJ.

**Fig. 2.**
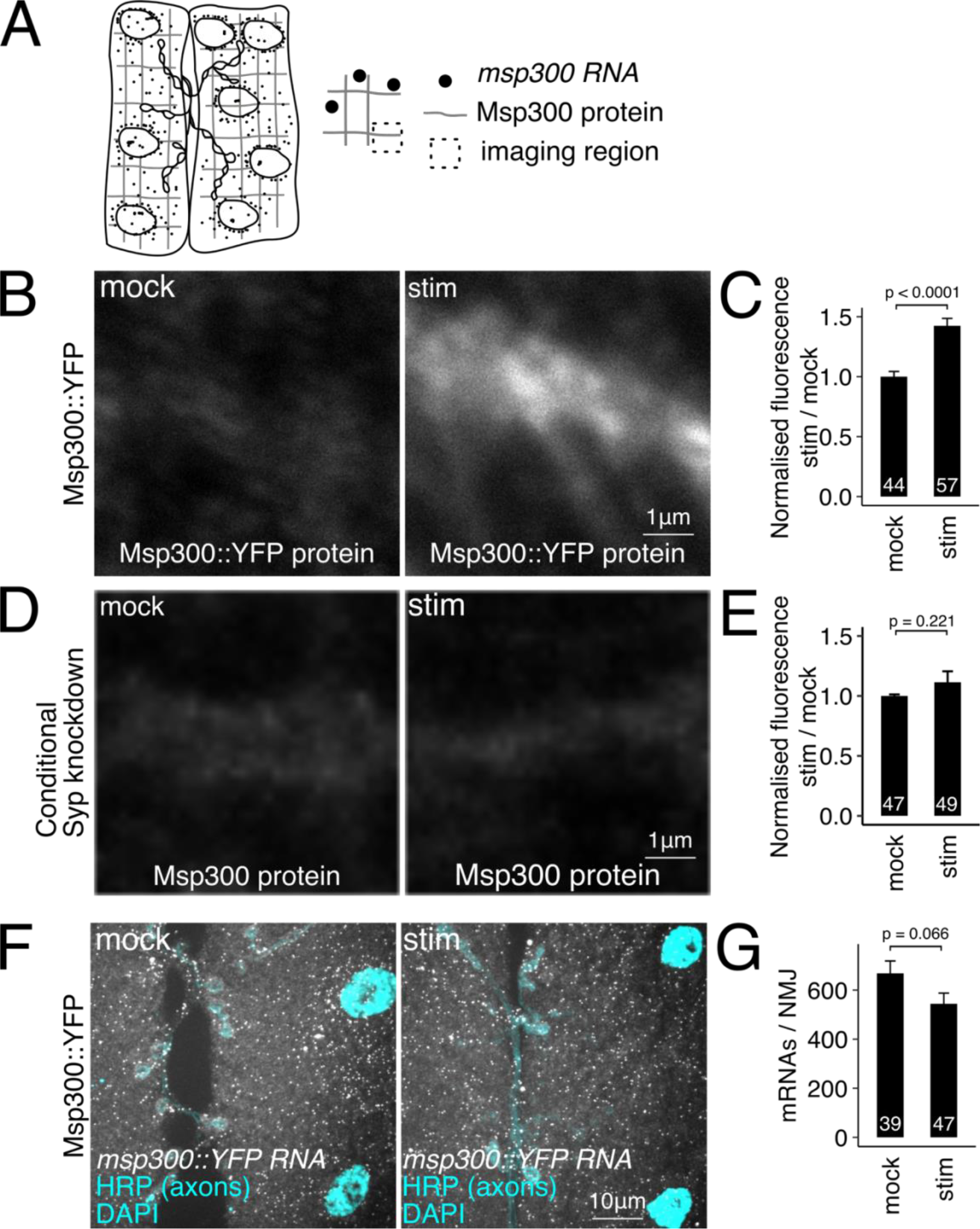
Activity-induced Msp300 expression requires Syp, and does not require *de novo* transcription. (A) Schematic showing the region (dotted square) that was imaged for immunofluorescence quantification. (B-C) Msp300 protein levels in stimulated larval NMJs are increased 40% relative to non-stimulated control NMJs. (D-E) Activity-dependent increase in Msp300 protein level is inhibited when *syp* expression is knocked down for 24hrs before the experiment. (F-G) Spaced potassium stimulation does not affect *msp300* mRNA levels at the NMJ. Mean ± sem; Student’s unpaired T-test; number of NMJs measured shown in each bar.

### Syp is required for activity-dependent plasticity at mature NMJ synapses

Syp was previously shown to be required for structural development of the synapse and synaptic transmission at the *Drosophila* larval NMJ (Halstead et al., 2014). We set out to test whether Syp is also required for activity-dependent synaptic plasticity by performing stimulus-induced plasticity assays in larval NMJs with *syp* knocked out or conditionally knocked down. Our results reveal a specific requirement for Syp in activity-dependent plasticity of mature synapses, in addition to synapse development. To assess activity-dependent plasticity at the NMJ we quantified new bouton formation and synaptic vesicle release in Syp mutants after KCl stimulation (Ataman et al., 2008). New bouton growth (ghost boutons; GBs) was quantified by counting the number of boutons labeled by HRP immunofluorescence that also lack the post-synaptic density marker Dlg1 (arrows, Fig. 3A). Wild type larvae produced an average of 4 GBs per NMJ in response to KCl, while mock-stimulated controls had an average of 0.5 GBs per NMJ. In the *syp* null mutant, *syp^e00286^*, the number of GBs induced by spaced KCl stimulation was two-fold less than wild type larvae (Fig. 3B). This phenotype is indeed due to loss of *syp* activity because it was also observed in hemizygous *syp^e00286^ /Df* NMJs, but not in P-element excision revertant larvae (Fig 3B). We conclude that Syp is required for *structural* plasticity at the larval NMJ.

**Fig. 3.**
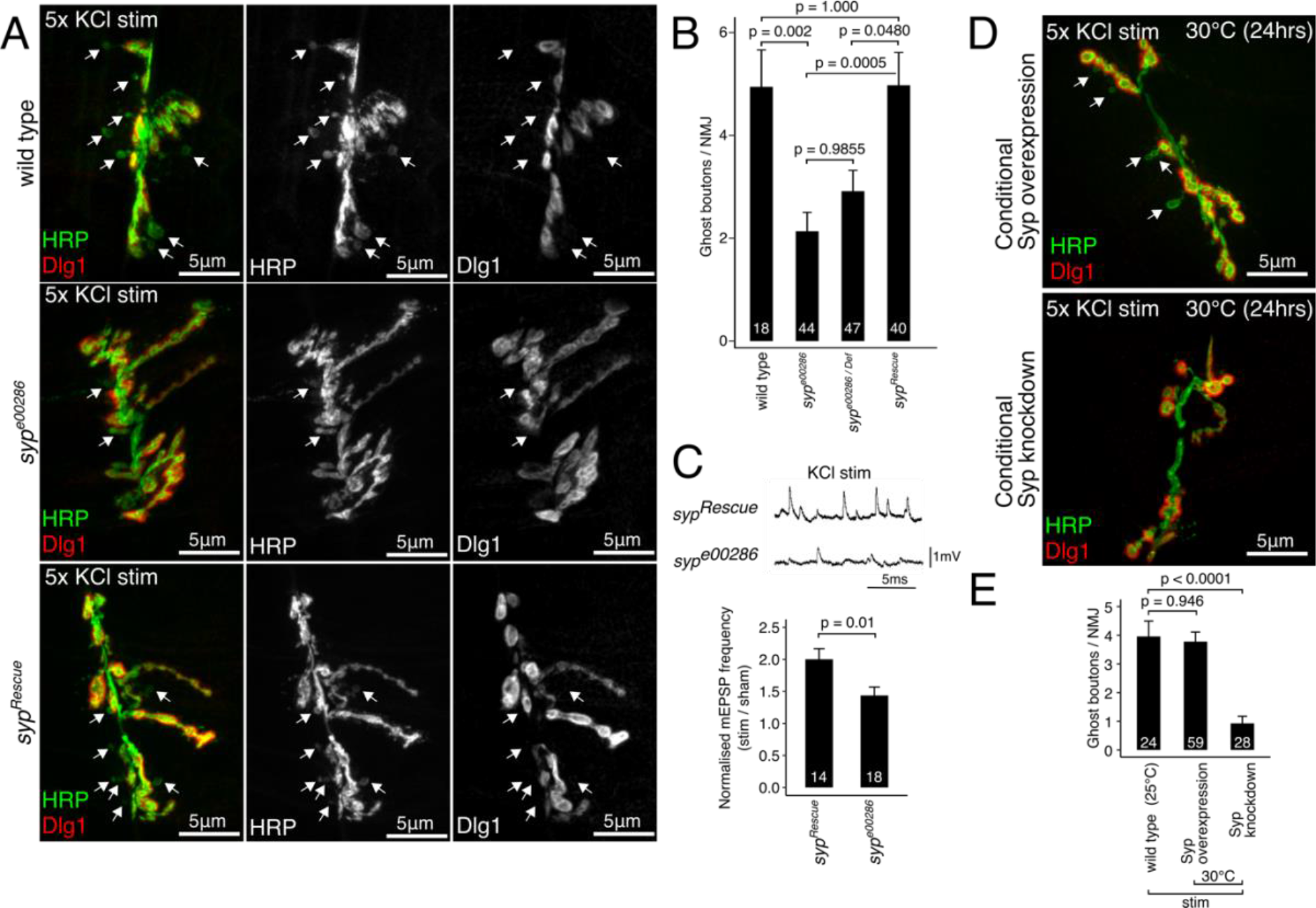
Syp modulates activity-dependent synaptic plasticity in the larval NMJ, both developmentally and at the mature synapses. (A) New synaptic boutons (ghost boutons, GBs; arrows) are formed by 5 rounds of KCl stimulation in wild type NMJ preparations. GBs appear as immature HRP-positive axon terminals (green) that lack the post synaptic density marker, Dlg1 (red). *syp* loss of function mutant (*syp^e00286^*) has abnormal synapse morphology and relatively few stimulus-induced GBs. (B) Quantification of stimulus-induced GBs per NMJ comparing wild type, *syp^e00286^*, *syp^e00286^/Def*, and P-element excision rescue larvae (mean ± SEM; Kruskal Wallis with Dunn’s post hoc test; number of NMJs shown in each bar). (C) Activity-induced potentiation of spontaneous synaptic vesicle release is inhibited in *syp* mutant larvae. Traces show mEPSPs recorded from muscles after KCl stimulus in *syp* mutant and *syp* rescue lines. Histogram shows the frequency of mEPSPs normalised to the average mEPSP frequency in mock-treated larvae (mean ± sem; Student’s unpaired T-test; number of muscles measured shown in each bar). (D) Larval NMJ morphology is unaffected by conditional overexpression or knockdown of *syp*. Representative max projection confocal images of NMJs from KCl stimulated larvae show the presence of GBs in the overexpression line, but not the conditional *syp* knockdown line. (E) Stimulus-induced GB formation is unaffected by conditional *syp* overexpression but conditional knockdown with *syp* RNAi completely abolished GB formation. Quantification of GB numbers (mean ± SEM; One-way ANOVA; number of NMJs is shown in each bar).

To assess whether Syp also has a role in *functional* activity-dependent synaptic plasticity we recorded miniature excitatory post-synaptic potentials (mEPSPs) after spaced KCl stimulation. Stimulus-induced potentiation of mEPSP frequency provides a physiological readout of NMJ plasticity (Ataman et al., 2008). In a Syp rescue line, the mEPSP frequency doubled in response to KCl stimulation, but was significantly less elevated in *syp*^*e00286*^ (Fig. 3C), indicating that Syp is involved in activity-induced potentiation of synaptic vesicle release. Together these results establish Syp as an important factor in structural and functional activity-dependent synaptic plasticity.

To separate the acute effects of Syp in synaptic plasticity from its developmental role in synapse formation we again used conditional knockdown of *syp* with Gal80^ts^/Gal4/UAS. By isolating Syp’s role at the mature NMJ from developmental effects, we were able to show that Syp acts immediately in response to neuronal activation to facilitate synaptic plasticity. Importantly, the NMJ morphology in conditional *syp* knockdown mutants was indistinguishable from wild type NMJs (Fig. 3D), indicating that synapse development was normal. Conditional *syp* overexpression did not interfere with GB formation in response to spaced KCl stimulation (Fig. 3E), however conditional *syp* knockdown completely inhibited KCl-induced GB formation (Fig. 3E). These experiments demonstrate a post-embryonic requirement for Syp in activity-dependent plasticity at mature NMJ synapses.

### Genetic, biochemical, microscopic, and biophysical evidence shows that Syp interacts with Msp300 at the larval NMJ

Having found that Syp is required for activity-dependent Msp300 expression and that both proteins are required for synaptic plasticity, we performed a series of experiments to determine whether *syp* and *msp300* directly interact. First, we determined that there is a genetic interaction between *syp* and *msp300*. Both are recessive genes such that activity-induced plasticity is normal in heterozygous larvae. However, activity-induced GB formation is completely abolished in larvae that are trans-heterozygous for both *syp* and *msp300* null alleles (Fig. 4A-E). These results demonstrate a functional link between *syp* and *msp300* that is specific to activity-dependent synaptic plasticity.

**Fig. 4.**
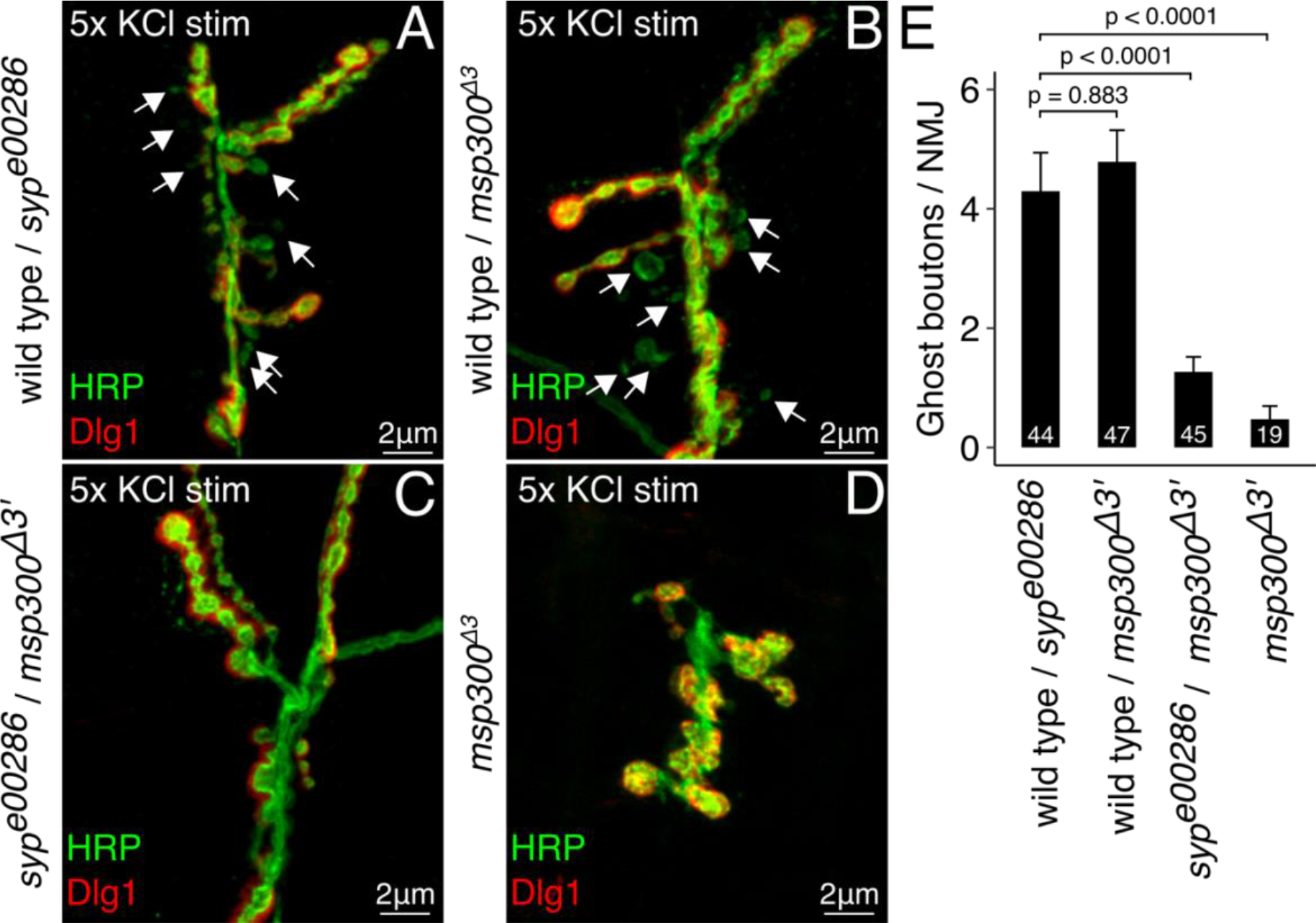
*msp300* and *syp* show a strong genetic interaction in activity-dependent bouton formation. (A-D) Trans-heterozygous *syp^e00286^*/*msp300*^*∆3’*^ mutant NMJs have normal bouton morphology but fail to produce ghost boutons (GBs) in response to KCl stimulation. Representative max projection confocal images from KCl stimulated NMJs show the presence of GBs in heterozygous *syp*^*e00286*^ (A) and *dNesp1*^*∆3’*^ (B) mutants, but not the trans-heterozygous *syp^e00286^*/*msp300*^*∆3’*^ mutants (C). (D) Homozygous *msp300*^*∆3’*^ mutants have significantly underdeveloped axon terminals and fail to produce GBs. (E) Quantification of KCl induced GB formation shows that *syp^e00286^*/*dNesp1*^*∆3’*^ mutants have significantly fewer activity induced ghost boutons than heterozygous controls (mean ± SEM; One-way ANOVA; number of NMJs is shown in each bar). Homozygous *msp300*^*∆3’*^ mutants and *syp*^*e00286*^ (Fig 1B) also exhibit the inhibited GB phenotype.

Next, we tested for biochemical interactions between Syp protein and *msp300* mRNA at the larval NMJ. Syp and *msp300* are both expressed post-synaptically at the NMJ, so we prepared lysates from dissected larval fillets after removing all other internal organs and the central nervous system. The presence of *msp300* mRNA in Syp immunoprecipitates was then quantified using RT-qPCR. Enrichment of m*sp300* mRNA in the IP fractions was on average 50-fold higher than the non-binding control transcript *rp49* (Fig. 5A), indicating that Syp associates with *msp300* transcripts in larval muscle, consistent with RIP-qPCR experiments from whole larvae (McDermott et al., 2014).

**Fig. 5.**
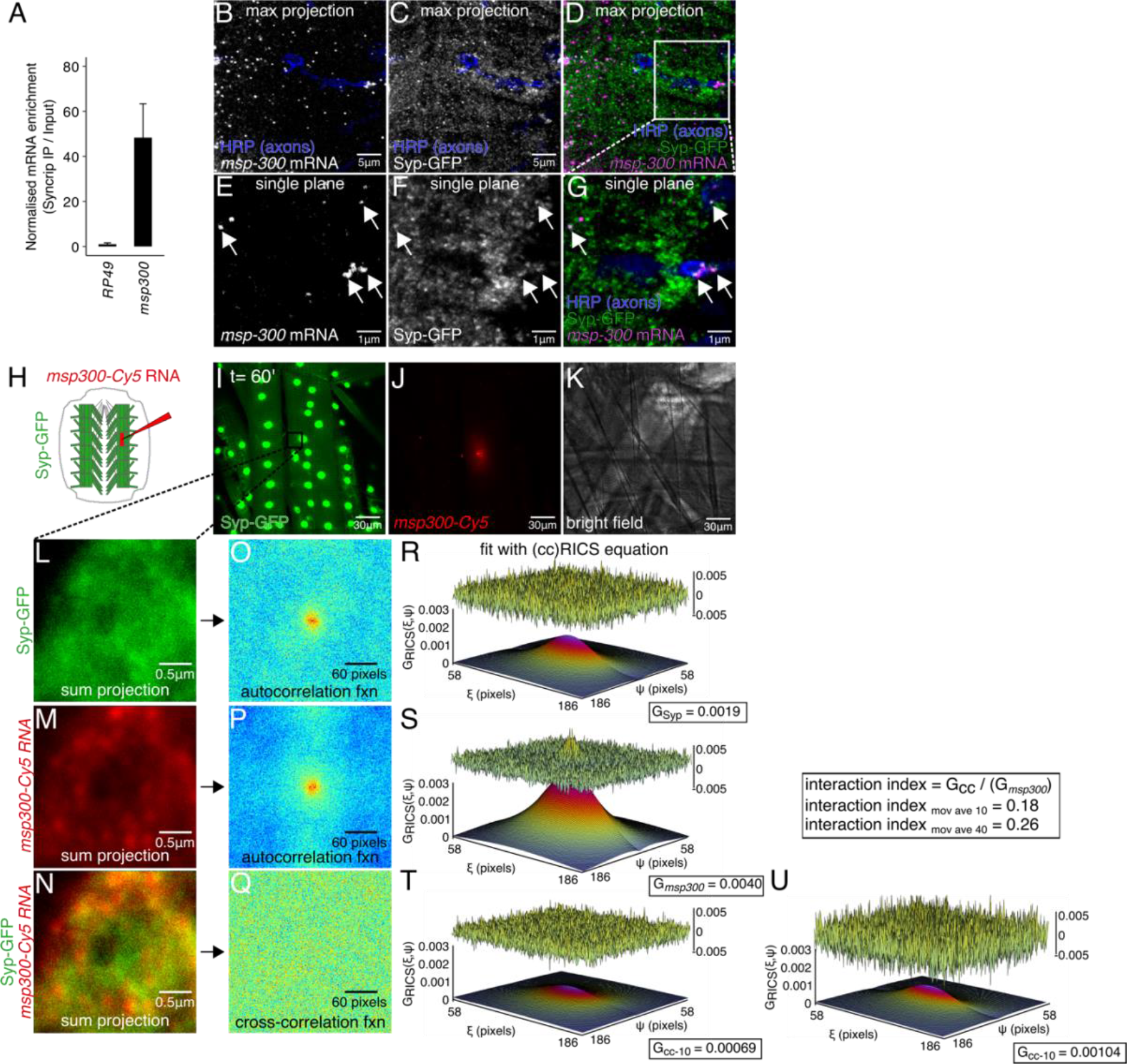
*msp300* mRNA physically interacts with Syp granules near the larval NMJ, *in vivo*. (A) *msp300* mRNA co-precipitates with Syp. Quantification of RT-qPCR data shows high enrichment of *msp300* relative to the non-binding control *rp49* (mean ± sem, N=3 IPs). (B-D) Representative confocal microscopy image of *msp300* smFISH and Syp-GFP signal at the larval NMJ, max z projection. (E-G) A single confocal slice of the same image, showing that *msp300* transcripts co-localise with Syp containing RNP granules within the resolution limit of the system. (H) Schematic of Cy5-labeled *msp300* RNA injection experiment in larval NMJ preparation to test its association with Syp. (I-J) Representative live images of Syp-GFP in a larval muscle injected with Cy5-labelled *msp300* mRNA. (K) A bright field image of an injected muscle indicates that the muscle still appears healthy 60 minutes after the injection. (L-N) Region of interest (from box in I) where ccRICS data were acquired. Each image is a sum projection of 50 images acquired in photon-counting mode. (O-Q) Autocorrelation function acquired from the images in L-N. (R-T) 3D plots of the spatial autocorrelation function with the relative amplitude and residuals after fitting with the RICS equation showing that ~12% of molecular complexes contain both Syp protein and *msp300* mRNA.

For interactions between Syp and *msp300* to be functionally relevant to activity-dependent plasticity they should occur near synapses. Our experiments show that Syp granules around the synaptic boutons contain *msp300* mRNA. We used two different techniques to assess the interactions between *msp300* mRNA and Syp protein at the NMJ. First we visualised individual *msp300* transcripts and Syp-GFP fusion protein in fixed NMJs using super resolution confocal microscopy (Korobchevskaya et al., 2017). Syp was tagged with a N-terminal eGFP fusion protein at the endogenous locus (see Methods for details). Expression of the Syp-GFP reporter is highly enriched in the nucleus and is also found in discrete punctae throughout the cytoplasm and near the synapse, as previously reported (Fig. 5B-D; (Halstead et al., 2014)). To co-visualise Syp and *msp300* we hybridised Syp-GFP larval fillet preparations with smFISH probes targeting *msp300*. After correcting the images for chromatic aberrations, we observed several *msp300* mRNA molecules that spatially overlapped with Syp-GFP foci (Fig. 5E-G, arrows), both in the cytoplasm and adjacent to the synapse. The presence of *msp300* transcripts in Syp granules near the NMJ suggests that local translation of *msp300* could be regulated by Syp.

To directly test for physical interactions between Syp and *msp300 in vivo*, we co-visualised *msp300* RNA and Syp protein in living larval NMJ preparations. We used cross correlation Raster Imaging Correlation Spectroscopy (ccRICS), a biophysical method that measures fluorescent protein complexes by virtue of correlated mobilities within an illuminated small rapidly scanned field (Digman et al., 2009). We microinjected Cy5 *in vitro* labelled *msp300* into the muscle cytoplasm of NMJ preparations from lines expressing endogenous Syp-GFP (Fig. 5H-K). The interaction between *msp300* RNA molecules and Syp-GFP molecules was quantified by measuring the fraction of *msp300* molecules that interact with Syp complexes. We found that on average, ~39% of *msp300* molecules interact with Syp-GFP at the larval NMJ. With ccRICS we were also able to determine that the interactions between RNA and protein are dynamic, as opposed to permanent association (see Methods for details).

As a negative control to test whether the interaction between Syp-GFP and *msp300* RNA was due to specific binding and not a random interaction between *msp300* RNA and the GFP fusion protein, we injected fluorescent *msp300* RNA into muscles expressing free GFP and acquired ccRICS data. We measured a negative interaction between *msp300* and free GFP, which means that ~3% of *msp300* RNA molecules are anti-correlated with cytoplasmic GFP, in contrast to ~39% positive correlation in the case of Syp-GFP (above). We conclude that *msp300* RNA dynamically interacts with Syp in RNP complexes at living NMJ synapses.

### Syp expression and protein dynamics are modulated by neural activity

Given Syp’s role in mediating activity-dependent gene expression changes, we next asked how Syp granules respond to elevated neuronal activity. We tested whether neuronal activation alters Syp abundance or diffusion rate using RICS. Our results show that KCl stimulation causes an increase in Syp abundance and a significant decrease in its diffusion rate. Syp-GFP fluorescence was measured in live NMJ preparations that were either stimulated with KCl or mock treated with normal HL3 saline, as a control. Images were acquired using the RICS format to measure Syp protein dynamics and Syp protein abundance from the same dataset (Fig. 6A). We found that Syp protein expression was elevated 1.6-fold in the cytosol (p < 0.0001; Fig. 6B-C) and 1.2-fold in the nucleus (p = 0.0274; Fig. 6B-C) of KCl stimulated samples relative to controls. From the same dataset we used RICS analysis to determine the diffusion rate of Syp *in vivo* (Fig. 6D-E). The average diffusion coefficient for Syp-GFP in mock stimulated samples was 0.84 ± 0.04μm^2^/s in the cytosol and 0.41 ± 0.03μm^2^/s in the nucleus respectively. In KCl stimulated samples there was a 25% decrease in Syp mobility relative to mock treated samples, specifically in the cytosol near the NMJ (Fig. 6F), suggesting that the size of Syp complexes is altered in the stimulated state.

**Fig. 6.**
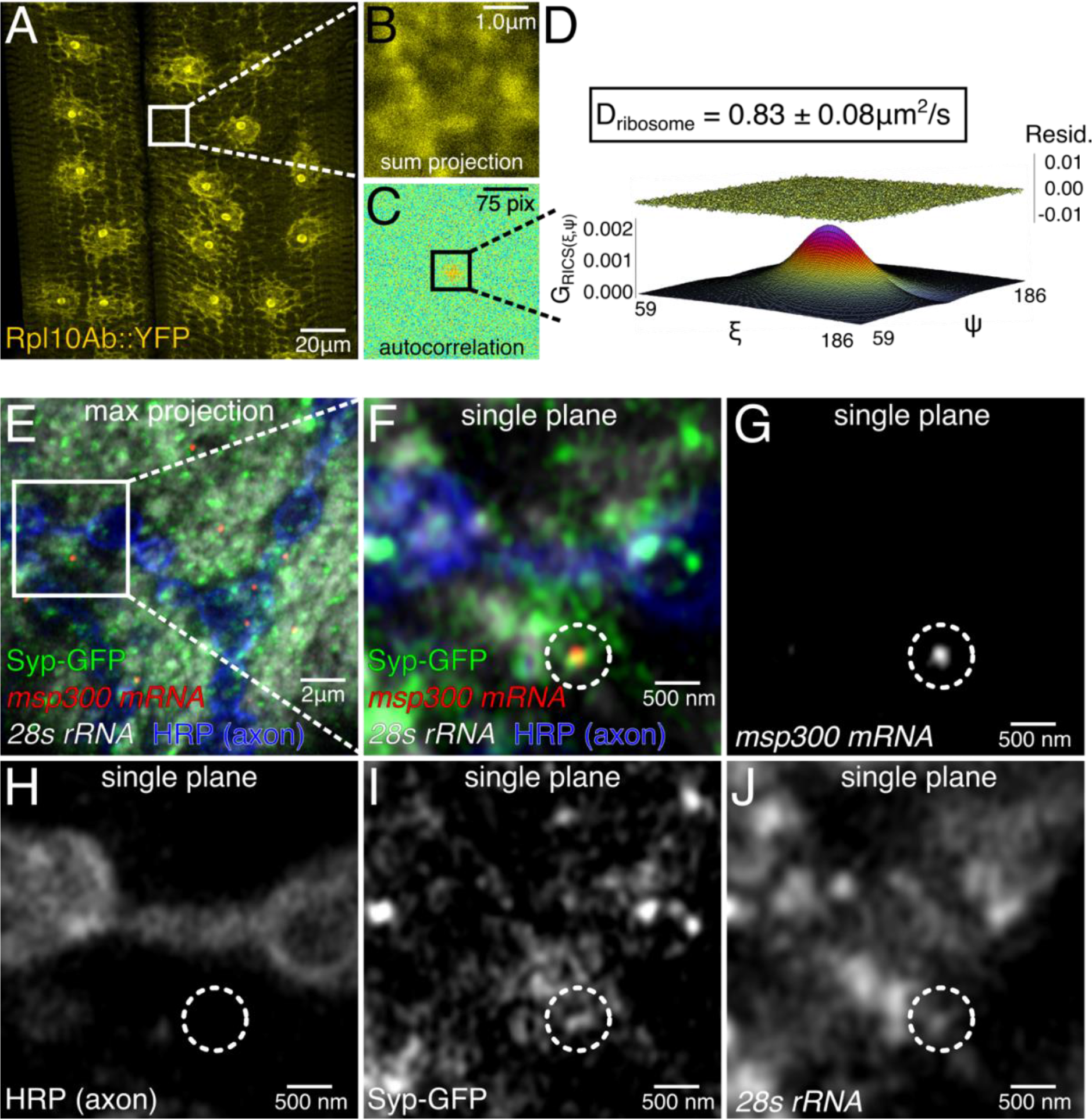
mRNP granules containing *msp300* mRNA and Syp protein are associated with ribosomes near the larval NMJ. (A-D) Diffusion rate of the large subunit ribosomal protein Rpl10A is almost identical to Syp. (A) Low magnification image of Rpl10A::YFP shows that the tagged protein is properly localised in muscle, i.e., in the nucleous and endoplasmic reticulum. Images taken in the RICS format (B) were used to calculate a spatial autocorrelation function (C) and determine the apparent diffusion coefficient (D). (E-J) Syp granules at the larval NMJ contain *msp300* mRNA and ribosomes. (E) Representative z-projection of Airyscan super resolution images showing Syp-GFP (green), *msp300* mRNA (red), and *28s ribosomal RNA* (white) at the synapse (blue). (F-J) Magnified single plane images show that an *msp300* transcript and rRNA reside within a Syp granule (dotted circle).

Activity-induced changes in the mobility of Syp could arise from a general increase in cytosolic viscosity. Similarly, the activity-induced increase in Syp protein levels could arise from a general increase in translation. To determine if activity-induced changes in Syp mobility and translation are specific, and not a general non-specific consequence of KCl stimulation on cellular viscosity and translation rate, we measured the effect of KCl stimulation on free GFP diffusion rate and protein expression levels (Fig. S4A,B). In mock-treated larval muscles the average GFP diffusion rate was 15.2 ± 4.7μm^2^/s, which is similar to GFP diffusion rates measured with various techniques in mammalian cells (Gura Sadovsky et al., 2017). The GFP diffusion rate in larval muscles was not significantly altered by KCl stimulation (Fig. S4C). Similarly, the level of cytosolic GFP was not significantly altered by KCl stimulation (Fig. S4C). Therefore, our control experiment showed that elevated synaptic activity does not cause a general, non-specific increase in translation nor a general change in the intracellular environment that influences protein diffusion. We conclude that activity-induced changes in Syp mobility and protein expression are a specific response to elevated synaptic activity.

To estimate the likely size of Syp-GFP complexes in live NMJs we compared the measured diffusion rate of Syp-GFP to free GFP, as obtained in the measurements above. Based on size alone, the Syp-GFP fusion protein is approximately 3-fold larger than GFP, from the Stokes-Einstein equation we predict that the diffusion rate of free GFP would be 1.4-fold faster than Syp-GFP. The measured GFP diffusion rate in larval muscles was ~18-fold faster than Syp-GFP, which strongly suggests that Syp exists in a large molecular complex. In human cell culture we also observed an un-expectedly slow Syncrip-GFP diffusion rate that was ~8-fold slower than GFP (Fig. S4E-I). These results are consistent with the notion that Syncrip is present in a large RNA granule in association with ribosomes and other RNA binding proteins.

To test whether the Syp complexes are likely to include ribosomes, we measured the diffusion rates of ribosomes in the larval NMJ. We found that ribosomal diffusion rates near the NMJ are very similar to Syp diffusion rates. Ribosome diffusion was measured by acquiring RICS data from a protein trap line with YFP inserted into the ribosomal Rpl10Ab gene (Lowe et al., 2014). The protein expression pattern of the Rpl10Ab::YFP protein trap line overlaps almost completely with an smFISH probe that detects 28s rRNA (Fig. S5A), indicating that Rpl10Ab::YFP gets incorporated into ribosomes. RICS analysis revealed that the average diffusion rate of Rpl10Ab::YFP near the axon termini was ~0.83 ± 0.08 μm^2^/s (Fig. 7A-D), which is nearly identical to that of Syp-GFP diffusion rate, and similar to diffusion rates measured for the large ribosomal subunit in mouse embryonic fibroblasts (Katz et al., 2016). These results suggest that Syp at the larval NMJ are both found in extremely large, similarly-sized complexes.

**Fig. 7.**
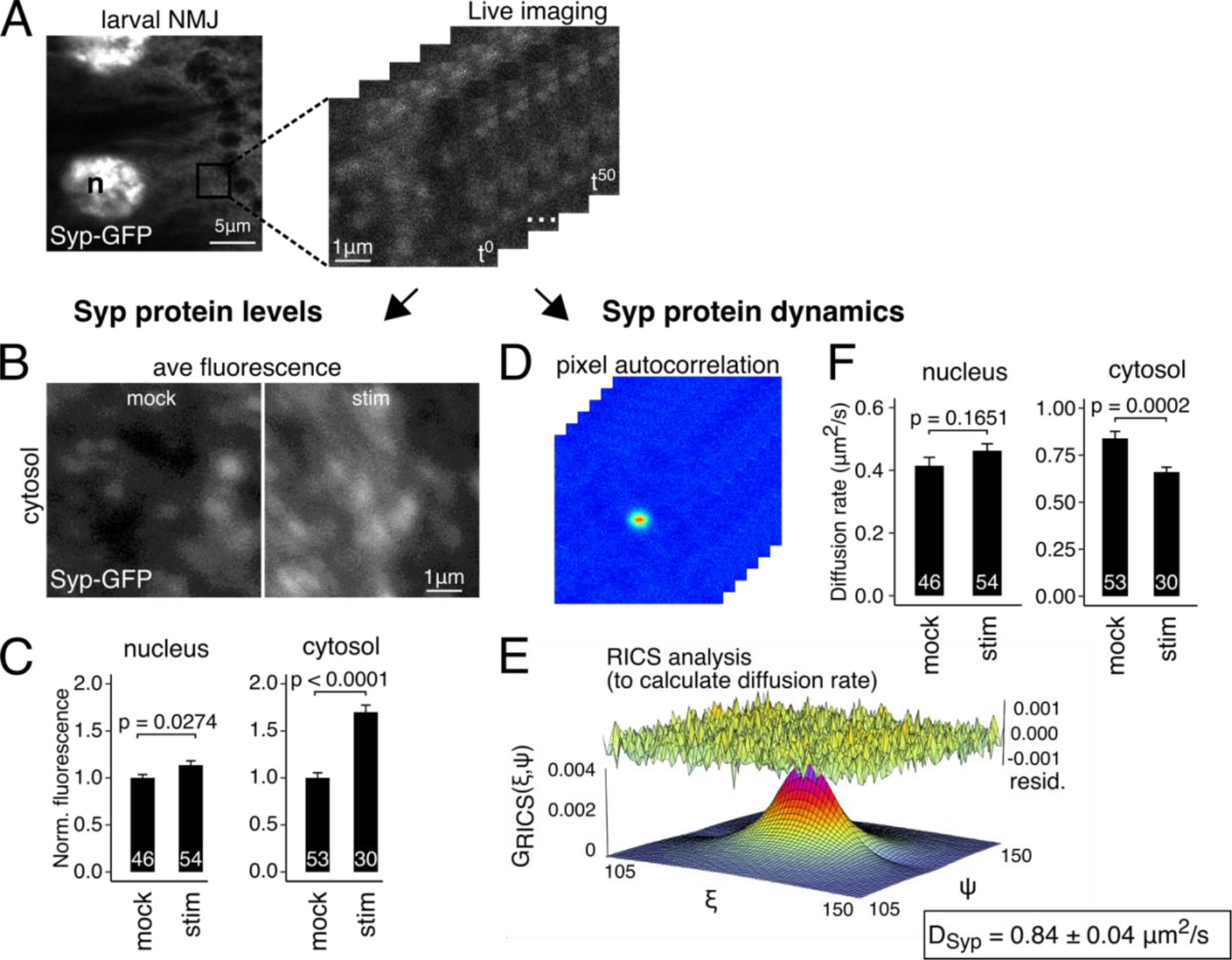
Synaptic activity modulates Syp protein levels and dynamics at the larval NMJ. (A) A series of scanning confocal images were acquired in raster imaging correlation spectroscopy (RICS) format to measure Syp protein levels and dynamics. (B) Average intensity projection of an image time series shows that Syp-GFP levels at the NMJ are higher in KCl-stimulated samples relative to mock-stimulated controls. (C) Quantification of fluorescence intensity shows that Syp-GFP levels in stimulated samples are significantly higher in the cytosol, but not in the nucleus. (D) A plot of the spatial autocorrelation function for each image after averaging across the time series and subtracting the immobile fraction. (E) 3D plot of the autocorrelation function fitted with RICS. RICS analysis was used to calculate the apparent diffusion coefficient for Syp-GFP in the measured regions. (F) Syp-GFP diffusion rate was significantly reduced at the NMJ in KCl-stimulated samples relative to mock-stimulated controls. Nuclear Syp-GFP diffusion was unaffected. Mean ± sem; Student’s unpaired T-test; number of NMJs measured shown in each bar.

### *msp300* mRNA molecules are localized at the larval NMJ in ribosome-containing Syp granules

To determine more directly whether Syp granules at the NMJ contain ribosomes we first used super resolution microscopy to image Syp-GFP together with *28s rRNA* smFISH in order to establish whether they are in the same sub-region of the cell. We found that Syp RNPs co-localise with *28s rRNA* and *msp300* mRNA, suggesting that ribosomes are present very near Syp RNP granules. We were not able to resolve the *28s rRNA* molecules as individual puncta because ribosomes are present at such a high concentration in the larval muscle (Zhan et al., 2016). In order to overcome this difficulty, we tried multiple super resolution microscopy techniques, including Airyscan, 3D structured illumination microscopy (3D-SIM), and STED. Though none of the techniques could resolve individual ribosomes, which measure less than ~30nm in their longest axis (Scofield and Chooi, 1982; Verschoor et al., 1996; Verschoor et al., 1998), we achieved x,y resolution between 90-150nm with the different techniques (Fig. S5B-P). This resolution allowed us to observe discrete Syp-GFP particles overlapping with *28s rRNA* particles in the sub-synaptic reticulum that resemble clusters of ribosomes described in electron micrographs of the NMJ (Fig. 7E-J) (Ukken et al., 2016; Zhan et al., 2016). We interpret these results to mean that close proximity with ribosomes and *msp300* mRNA allows Syp to rapidly facilitate translation at the synapse in response to elevated neural activity.

## Discussion

Neuronal synapses form through a genetically determined developmental program followed by continuous adaptation to new neuronal stimuli. This process is called synaptic plasticity and is characterized by building new synapses and changing the molecular composition of existing synapses (McAllister, 2007; Munno David and Syed Naweed, 2004). Synaptic plasticity forms the basis of memory and learning and is perturbed in numerous neurological diseases including neurodegeneration. There has been considerable progress in identifying gene expression changes that underpin synaptic plasticity, with a predominant focus on transcriptional regulation that is critical during development and learning. Much less is known about the role of post-transcriptional mechanisms in plasticity, despite the well-recognized problem that transcriptional regulation alone cannot explain all of the processes of activity-dependent synaptic plasticity. Immediate activity-dependent gene expression changes are too rapid to be explained by new transcription alone and instead require mechanisms that can act near the synapse, a long distance from the cell nucleus. Moreover, the local changes in the actin cytoskeleton that are associated with new synapse formation require changes in the levels of actin regulating proteins to be controlled locally. Here, we investigated local expression of the actin regulating protein Msp300, one of the first proteins to accumulate at new synaptic boutons, and which is essential for new synapse formation. By identifying the RBP Syp as a regulator of Msp300 expression, we have discovered an important post-transcriptional mechanism in activity-induced gene expression and synapse formation. These findings provide a new mechanism to explain rapid local actin reorganization near the synapse, which is essential for new synapse formation.

We performed a series of biochemical, biophysical, and imaging experiments to determine that *msp300* and Syp interact at the larval NMJ, *in vivo*. Our results also show that the interactions between *msp300* and Syp are important for activity-dependent synaptic plasticity, as we observed a strong genetic interaction between *msp300* and *syp* that inhibited activity-induced bouton formation (Fig. 4E), and mutants with single homozygous mutations in either *syp* or *ms300* also exhibit the same phenotype (Figs. 3B,4E). We find that Syp RNP granules contain *msp300 mRNA* ribosomes (Fig. 6), which suggests that they are translationally competent. Finally, the mobility of Syp granules changes in response to stimulation, which can be explained by an increase in ribosome density in the granule as a consequence of an increased rate of translation. This idea is compatible with previous reports showing that Syp influences the level of proteins protein produced from its mRNA target, *grk*, in the *Drosophila* oocyte (McDermott et al., 2012). Syp has also been shown to facilitate translation in a wide range of biological contexts, including HIV-1 Gag-p24 RNA (Vincendeau et al., 2013), Hepatitis C viral mRNA (Kim et al., 2004), the circadian clock gene Per1 (Lee et al., 2012), and the p53 tumor suppressor (Kim et al., 2013). While we do not directly demonstrate translation of *msp300* in mRNA granules, taking our results in the context of the existing literature strongly suggests that Syp acts directly on *msp300* at the level of translation.

Our data support a model in which the rapid enrichment of Msp300 at new synapses is achieved by translation of localised *msp300* transcripts that are contained in Syp granules (Fig. 8). We show that Syp granules at the NMJ become significantly less dynamic after KCl stimulation (Fig. 7), which either means that the size of the complex increases or transport of the complex decreases. Our data indicate that interactions between Syp and *msp300* are largely through dynamic binding events rather than co-diffusion in a stable complex, which we interpret to mean that activity-induced translation occurs when localized *msp300* transcripts interact with Syp granules that have recruited additional ribosomes and translation machinery. The frequency of interactions should increase as a consequence of the 60% increase in Syp expression induced by the stimulus (Fig. 7B,C), thereby increasing local translation of *msp300* transcripts to accumulate Msp300 protein at the synapse and organise the actin scaffold. Although our data do not rule out the possibility that existing Msp300 protein is redistributed to new synapses upon activation, we favour the hypothesis that Syp facilitates activity-induced translation of *msp300* at the synapse by enabling the recruitment of ribosomes and translation factors to *msp300* mRNA.

**Fig. 8.**
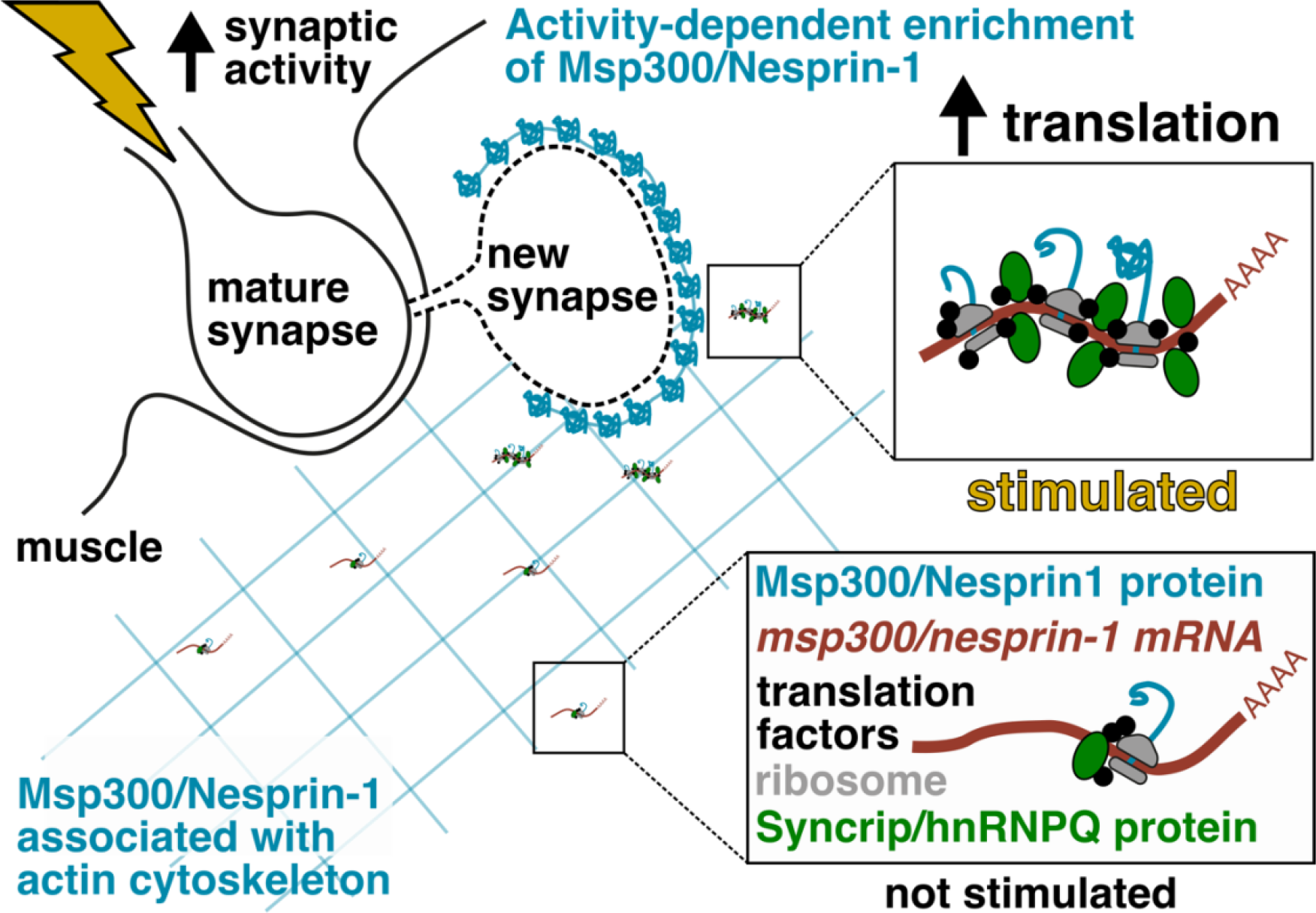
Proposed mechanism for Syp’s role in regulating activity-dependent enrichment of Msp300 and new synapse formation. Synaptic activity at the larval NMJ is elevated by increased crawling behaviour in the animal, which induces the formation of a new synaptic bouton. Msp300 is rapidly enriched around the new bouton to organize an actin scaffold where post synaptic proteins will be anchored. *msp300* mRNAs at the synapse are in a ribosome-containing complex with Syp that becomes much less dynamic in response to elevated activity. We hypothesise that Syp facilitates activity-dependent translation of *msp300* at the synapse by enabling recruitment of additional ribosomes and translation factors to the mRNA.

*Drosophila msp300* is orthologous to the mammalian genes *SYNE-1* and *SYNE-2*. *msp300* and *SYNE-1/2* both encode Nesprin proteins that perform many similar functions, including regulation of glutamate receptor expression (Cottrell et al., 2004; Morel et al., 2014) and positioning of myonuclei (Stroud et al., 2017; Volk, 2013; Wang et al., 2015; Zhou et al., 2018a). The functional role of Nesprins in the nervous system is more enigmatic and requires further attention because mutations in *SYNE-1* and *−2* are strongly linked to recessive forms of hereditary cerebellar and extra-cerebellar ataxias in humans (Dupre et al., 2007; Gros-Louis et al., 2007; Noreau et al., 2013; Synofzik et al., 2016; Wiethoff et al., 2016). The molecular function that causes central nervous system specific defects in these ataxias is not known, though it has been shown that neurogenesis and neuronal migration are significantly impaired (Zhang et al., 2009) and that white matter, cerebellar and cortical regions of the brain were significantly disrupted in patients (Gama et al., 2018). The *SYNE-1* ataxias also demonstrate extracellebellar phenotypes that are similar to neurodegenerative disease (Gama et al., 2016; Mademan et al., 2016). The most common mutations already observed to be linked with *SYNE-1* ataxias cause truncations or abnormal splice junctions. Our results suggest that impaired post-transcriptional regulation of *SYNE-1* could exacerbate the ataxia phenotypes.

Our work provides insight into how the actin cytoskeleton is regulated during activity-dependent synaptic plasticity. A filamentous actin scaffold is required at new synapses to anchor post-synaptic density proteins, which in turn anchor post-synaptic receptors. Msp300 is one of the first proteins to assemble at newly formed synaptic boutons (Fig. S1) where it is thought to facilitate actin polymerization by recruiting an unconventional myosin from the Myosin ID family, Myo31DF in *Drosophila* (Packard et al., 2015). Myo31DF forms a complex with Arp2/3 to mediate actin nucleation (Evangelista et al., 2000). How these actin regulating factors assemble at the synapse is not yet known, but local translation is an attractive mechanism since it is already well established that beta-actin is translated from localized mRNA in mammalian dendrites (Buxbaum et al., 2014; Eom et al., 2003; Katz et al., 2016). There is also extensive evidence showing that *arc1* mRNA, a cytoskeleton-associated immediate early gene, localizes to dendrites and is locally translated (Guzowski et al., 2000; Steward et al., 1998; Steward and Worley, 2001). Several additional mRNAs that encode actin regulating proteins, including Arp2/3, have been identified by sequencing transcriptomes specifically from dendritic compartments (Will et al., 2013), and Syp binds several mRNAs in addition to *msp300* that encode actin regulating proteins. Thus, more work is needed to determine if post-transcriptional regulation of localized actin regulating proteins, by Syp in particular, may be a general and conserved mechanism that is important for synaptic plasticity in various brain regions.

## Conclusion

We identified a key component of post-transcriptional regulation during activity-dependent synaptic plasticity at the larval NMJ, which provides insight into how an actin binding protein is locally enriched to organize the post-synaptic scaffold for new synaptic boutons. Syp and Msp300 are known to be present at various synapses in other organisms, so it is tempting to speculate that interactions between Syp and *msp300* transcripts will be required for synaptic plasticity in those systems. Our study also lays the groundwork for studying biophysical properties of mRNA and associated granules at intact synapses, *in vivo*. Extending this approach to other molecules will provide a more complete picture of how the cell consolidates experience into new synapses.

## Acknowledgements

This work was supported by a Wellcome Senior Research Fellowship (096144) and Wellcome Investigator Award (209412) to I.D. Advanced microscopy facilities and technical advice were provided by Micron (http://micronoxford.com), supported by Wellcome Strategic Awards (091911 and 107457) and an MRC/EPSRC/BBSRC next generation imaging award. D.I.-H. was supported by University College London. E.G. was supported by the National Institute of General Medical Sciences (NIGMS) NIH grant 2P41GM103540. F.R. was funded by a Marie Skłodowska-Curie Postdoctoral Fellowship. We are very grateful to the Bloomington *Drosophila* Stock Centre (Fly Stocks and cDNA) and to Flybase for their reagents and open data, which were both invaluable to this work. We would like to thank members of the Davis Lab for critical reading of the manuscript and Dr. Talila Volk for antibodies, stocks and discussions. We would also like to acknowledge Drs. Christoffer Lagerholm, Richard Parton, Lothar Schermelleh, and Pablo Hernandez-Varas for assistance and advice on super resolution microscopy and specimen preparation.

## Author contributions

JT, D.I.-H., and ID conceived and designed the study, interpreted the results and wrote and revised the manuscript.

JT Performed the majority of the experiments, reagent generation and data analysis. FR, and AJ generated additional reagents and performed additional experiments.

CS, and EG performed additional data analysis.

## Methods

### *Drosophila melanogaster* maintenance

Fly stocks were maintained with standard cornmeal food at 25ºC on 12hr light:dark cycles unless otherwise specified. Wandering third instar larvae were used for all experiments. The following genotypes were used: Oregon R (wild type unless otherwise specified), *syp*^*e00286*^ (McDermott et al., 2014); MHC-Gal4 muscle driver, UAS-Syp-GFP (McDermott et al., 2014), C57-Gal4 muscle driver, tubulin-Gal80^ts^, and the following MS2/MS2 coat protein (MCP) lines: hsp83-MCP-mCherry (Hayashi et al., 2014), hsp83-MCP-GFP and *grk*-MS2×12 (described in: Jaramillo et al., 2008). The UAS-Syp RNAi line was obtained from Vienna Drosophila Stock centre and was previously characterized in the larval NMJ (McDermott et al., 2014). All other lines were obtained from Bloomington Drosophila Stock Centre.

### Whole mount single molecule *in situ* fluorescence hybridisation (smFISH) and immunofluorescence (IF)

Stimulated larval NMJ specimens or mock treated controls were prepared using a protocol that has been described previously (Titlow et al., 2018). Briefly, specimens were fixed in paraformaldehyde (4% in PBS with 0.3% Triton-X; PBTX) for 25mins, rinsed three times in PBTX, blocked for 30min in PBTX+BSA (1%), and incubated overnight at 37°C in hybe solution (2x SSC, 10% formamide, 10% dextran-sulfate, smFISH probes [250nm], and primary antibodies). The next day samples were rinsed 3x in smFISH wash buffer (2x SSC + 10% formamide) and incubated for 45min at 37°C in smFISH wash buffer with secondary antibodies and DAPI, then washed for 30min in smFISH wash buffer at room temperature before mounting in glycerol (Vectashield). PBTX was used in place of smFISH buffer for experiments that did not require smFISH. The following antibody (concentrations) were used: mouse anti-Dlg1 (1:500), guinea pig anti-Syp (1:500), HRP-Dyelight-405/488/Alexafluor-568/Alexfluor-659 (1:100), Donkey anti-guinea pig-Alexafluor-488 (1:500), donkey anti-mouse Alexafluor-568 (1:500).

### Image acquisition and analysis

Whole-mounted immunofluorescence and smFISH specimens were imaged on a spinning disk confocal (Ultra-View VoX-PerkinElmer) with 60x oil objective (1.35 NA, UPlan SApo, Olympus) and emCCD camera (ImagEM; Hamamatsu Photonics). NMJs at muscles 6 and 7 in segments 3-5 were imaged for at least five different larvae per condition/genotype. Ghost boutons (GBs) were counted manually. Immunofluorescence signal intensity was quantified by measuring the average of 10 small regions (1×1μm square) from average intensity z-projections of each NMJ. Mature and nascent transcripts were counted using a Matlab program called FISHquant (Mueller et al., 2013).

Super resolution images were acquired on an LSM-880 (Zeiss) with Airyscan detector and 60x/1.4NA oil objective. Main pinhole was adjusted to 2.0AU with 0.2AU pinhole in each of the 32 individual channels. Voxel size was set to 40nm in x,y, 150nm in z. To correct for chromatic aberration, we labelled the DNA with Vybrant® DyeCycle™ Violet Stain (which emits from blue to far-red spectra) and acquired z-stacks in each emission channel with 405nm excitation. Chromagnon (Matsuda et al., 2018) was then used to apply chromatic shift correction to images where co-localisation was assessed.

To assess the performance of Airyscan relative to other super resolution microscopy techniques we acquired images of *28s rRNA* smFISH at the larval NMJ using SIM and STED microscopy. The x,y spatial resolution of each modality was then estimated from fast Fourier transform radial plots generated in SimCheck (Fig. S6G-O) (Ball et al., 2015). SIM images were acquired on a DeltaVision OMX V3 (GE Healthcare) with 60x/1.42NA oil objective (PLAPON 60XO NA1.42; Olympus) and Cascade II 512 EMCCD cameras (Photometrics). SIM reconstruction was performed with softWoRx (GE). STED images were acquired on a Leica TCS SP8 STED 3X inverted microscope with HC PL APO 93X/1.30NA glycerol objective and GaAsP HyD detector.

### Spaced potassium stimulation protocol

Six third instar larvae were dissected in two separate chambers to allow even saline perfusion from peristaltic pumps. Five applications of high potassium saline (KCl) were separated by 15min perfusion of HL3 as described previously (Attaman et al., 2009). For smFISH and IF the larvae were fixed 150min after the first stimulation. For electrophysiology and live imaging experiments the recordings were made from 10min after the last stimulus.

### Electrophysiology

Groups of three larvae were analysed after chemical activation or mock treatments. Intracellular recordings were made in muscle 6 in segments 3-5 using sharp glass electrodes (10-20MΩ) filled with 3M KCl. Miniature EPSPs were amplified with a Multiclamp 200B, digitised with a Digidata 1550A A/D board controlled with pClamp (v10, Molecular Devices), and analysed offline using Mini Analysis software (v6.0.3, Synaptosoft). Spontaneous activity was recorded for two minutes and mEPSP frequency was analysed for the second minute.

### RNA immunoprecipitation and RT-qPCR

For each biological replicate, 10 third instar larval body walls were dissected in HL3 medium and homogenised in immunoprecipitation (IP) buffer (50 mM Tris-HCl pH 8.0, 150 mM NaCl, 0.5% NP-40, 10% glycerol, 1 mini tablet of Complete EDTA-free protease inhibitor (Roche) and RNAsin (Promega)). Lysates were incubated overnight at 4°C with magnetic Dynabeads (Thermo Fisher Scientific) conjugated to Guinea pig anti-Syp and IgG antibody. Beads were washed four times briefly with cold lysis buffer. To retrieve the RNA, beads were re-suspended in elution buffer (50 mM Tris-HCl pH 8.0, 10 mM EDTA and 1.3% SDS, RNAsin) and incubated at 65 °C, 1000 rpm for 30 min on thermomixer. The elution step was repeated and the supernatants were pooled. RNA was then extracted using an illustra RNAspin mini kit (GE healthcare). Input and eluate samples were used for cDNA synthesis using RevertAid Premium Reverse Transcriptase (Thermo Fisher Scientific). cDNA was used directly as a template for real-time PCR (SYBR green, Bio Rad).

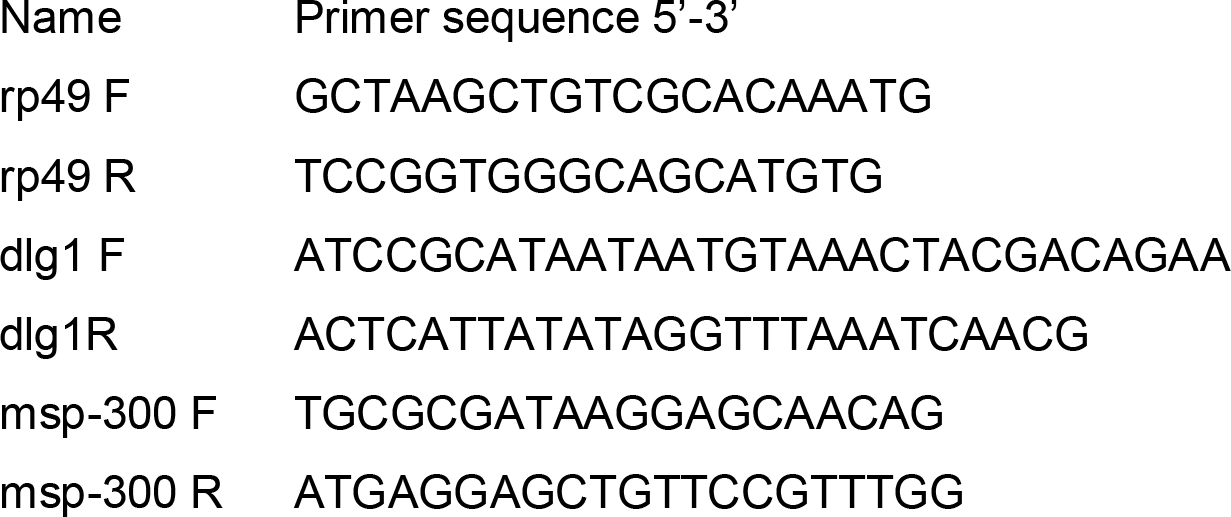

### Generation of Syp-GFP fly line

The syp-AttP line was generated using CRISPR to delete a 4Kb section at the beginning of the syp coding region, which was replaced by an AttP site. sgRNA construct design and validation was performed by Dr. Andrew Basset - Genome Engineering Oxford (GEO). 1kb Homology arms, corresponding to sequences flanking the sgRNA cleavage sites (located in the 5’UTR and 3rd intron of isoform F), were cloned into the pDsRed-attP vector (Addgene 51019). sgRNA constructs and the homology construct were injected in vas-cas9 embryos (BL 51323) by the Cambridge Fly facility. Embryos from the syp-AttP line were then injected with an AttB construct (RIV^Cherry^, Baena-Lopez et al., 2013) containing eGFP fused to the N terminus of Syp.

### sgRNA Guide sites

sgRNA5’ TGCGTTCGTTGAACTCTACAAGG

sgRNA3’CCTTTCGATTTGGGGGGATATGG

### Doxycycline-induced expression of GFP and Syncrip-GFP in HeLa cell lines

To generate stable cell lines, eGFP and human Syncrip-GFP plasmids were cloned into the Flp-In™ expression vector and integrated into Flp-In™ 293 T-REx cells using standard procedures. Briefly, we obtained the plasmids from ____… For RICS imaging experiments, the cells were grown to 50% confluence in duplicate cultures on 6-well plates (9cm^2^) and induced with doxycycline (0.10 μg/mL in clear DMEM with 10% BSA) for six hours prior to imaging. Cells were then imaged in a temperature-controlled chamber at 37°C.

### Raster imaging correlation spectroscopy (RICS) and cross correlation RICS (ccRICS)

The RICS method derives the apparent molecular diffusion rate from calculation of the spatial autocorrelation function between points in a scanning confocal image (Brown et al., 2008; Digman et al., 2009; Rossow et al., 2010). RICS data were acquired on a Zeiss LSM-880 upright confocal system using a 20x/1.0NA water immersion objective (Plan Apo; Olympus) and GaSP detector in photon counting mode. Laser power, pixel dwell time (8.19μs), pixel size (20nm), and frame size (256×256 pixels) were kept constant for all specimens and mock-treated specimens were always measured in parallel with stimulated specimens. Fluorescence intensity was quantified as average raw pixel intensity values from an average of 50 individual frames.

Calibration data were acquired as described above for live imaging, i.e., 50 frames (256×256) were acquired with constant pixel width (20nm), pixel dwell time (8.19μs), and line scan time (4.92ms). Calibration images were performed each day to determine the size of the beam waste. Donkey IgG conjugated to Alexa488 (10nM) was imaged with the settings described above and the data were fitted in SimFCS software (Brown et al., 2008) using 40μm^2^/s as the defined value for diffusion coefficient (Arrio-Dupont et al., 2000). Moving average of 10 frames was applied to remove artefacts from cell movement and the data were fit with a single component model, as residual plots and chi-square values revealed an acceptable goodness of fit.

ccRICS was used to calculate the proportion of fluorescent *msp300* RNA molecules interacting with Syp-GFP complexes. Images of both molecules were acquired simultaneously from two channels, corrected for chromatic shift as described above (using 100nm Tetraspek beads instead of Violet Dye) and fit with RICS and ccRICS equations. The interaction index was calculated by dividing the amplitude of the ccRICS autocorrelation function (G_cc_) by the amplitude of the RICS autocorrelation function of the *msp300* channel (G_*msp300*_). The result is an interaction index (G_cc_ /G_*msp300*_) that is proportional to the fraction of *msp300* molecules interacting with Syp complexes. We calculated the amplitude of the cross-correlation RICS function (G_cc_) using a moving average of 10 frames to subtract the immobile fraction, or a moving average of 40 frames to assess binding interactions (Digman et al., 2009). The interaction index (G_cc_ /G_*msp300*_) derived from a ccRICS function with a moving average of 10 frames was 0.13 ± 0.04, which means that ~13% of *msp300* mRNA molecules diffuse in a complex with Syp molecules. The interaction index was 0.39 ± 0.10 when derived from a ccRICS function with a moving average of 40 frames, which means that ~39% of *msp300* mRNA molecules interact transiently with Syp complexes. As a negative control, we measured the interaction index for *msp300-Cy5* RNA injected into larval muscles expressing cytosolic GFP. We found that *msp300-Cy5* was anti-correlated with cytosolic GFP, regardless of whether the moving average for the ccRICS function was 10 frames (interaction index = −0.002 ± 0.0005) or 40 frames (interaction index = −0.03 ± 0.03).

### Fluorescence correlation spectroscopy

Fluorescence correlation spectroscopy (FCS) data were acquired on a Zeiss LSM-880 upright confocal system using a 20x 1.0NA water immersion objective and GaASP detector with factory photon counter. Photon counts were acquired at 15MHz for 10s intervals at different spots throughout the muscle and the correlated data (1ms bins) were saved as .fcs files. The autocorrelation function was fit with the standard equation for 3D diffusion:

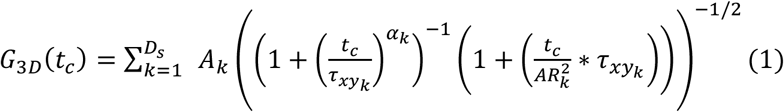

where G_3D_ is the amplitude of the correlation function, t_c_ represents time, D_s_ is the number of diffusing species, A_k_ is a factor that establishes the proportion of each diffusing species, τ_xy_ is the lateral diffusion rate, α is the diffusion anomalous diffusion factor, and AR is a constant factor that relates the axial diffusion rate to the lateral diffusion rate (Waithe et al., 2016). For GFP diffusion in the muscle the autocorrelation was fit with a 1 species diffusion model, anomalous diffusion factor of 1, and AR factor of 5.

### *In vitro* transcription and microinjection of Cy5-labelled *msp300* RNA

Msp300 template cDNA from the *Drosophila* Gold collection (HL01686; Rubin et al., 2000) was amplified by bacterial transformation using a standard protocol from the *Drosophila* Genomics Resource Center. The plasmid DNA (5μg) was linearised in an overnight digestion reaction with Apal and cleavage was verified by gel electrophoresis. After purifying the plasmid with QIAquick PCR Purification Kit, *msp300-Cy5* RNA was transcribed in a 50μL reaction according to the polymerase manufacturer’s instructions using the following components: T3 polymerase and associated transcription buffer (20units; ThermoScientific), linear DNA (1μg), mCAP analogue (Stratagene), DTT (1M), rNTP mix-UTP (10 mM CTP; 10 mM ATP; 3 mM GTP), Cy5-labelled UTP mix (1:1 mixture of labelled and un-labelled UTP, 10mM total [UTP]), and RNAse inhibitor (40units; Promega). Template DNA was digested with RNAse-free DNAse1 (2.0 units; Qiagen), and *msp300-Cy5* RNA was purified using a Sephadex G50 spin column (Roche miniQuick Spin RNA column), followed by EtOH precipitation. Purified RNA was then diluted to 100ng/μL with RNAse-free water for injection.

Cy5-labelled *msp300* RNA (Cy5-UTP) was pressure injected into *Drosophila* larval muscles using pre-fabricated glass capillary tips (0.5μm inner diameter, 1.0 μm outer diameter; Eppendorf Femptotips). Short pulses (3-5 x 100ms) were delivered into muscle 6, and delivery of the labelled RNA was verified by epifluorescence. Specimens were then transferred to a Zeiss LSM-880 scanning confocal for ccRICS analysis.

### Statistical analysis of ghost bouton, mEPSPs, mRNA, and protein levels

Statistical tests that were applied to each dataset are given in the Figure legends along with the number of samples appearing in each graph. The normality assumption was tested with the Shapiro-Wilk test. The equal variances assumption was tested with an F-test or Levene’s test, depending on the number of groups. Normally distributed populations with equal variances were compared using Student’s t-test or one-way ANOVA (with Tukey test for multiple comparisons), depending on the number of groups. Populations with non-normal distributions were compared using the Wilcoxin rank sum test or Kruskal-Wallis test (with Dunn test for multiple comparisons), depending on the number of groups. All statistical analyses were performed in R (Version 3.3.2 running in Jupyter Notebook).

## Supplementary Figures

**Figure S1.**
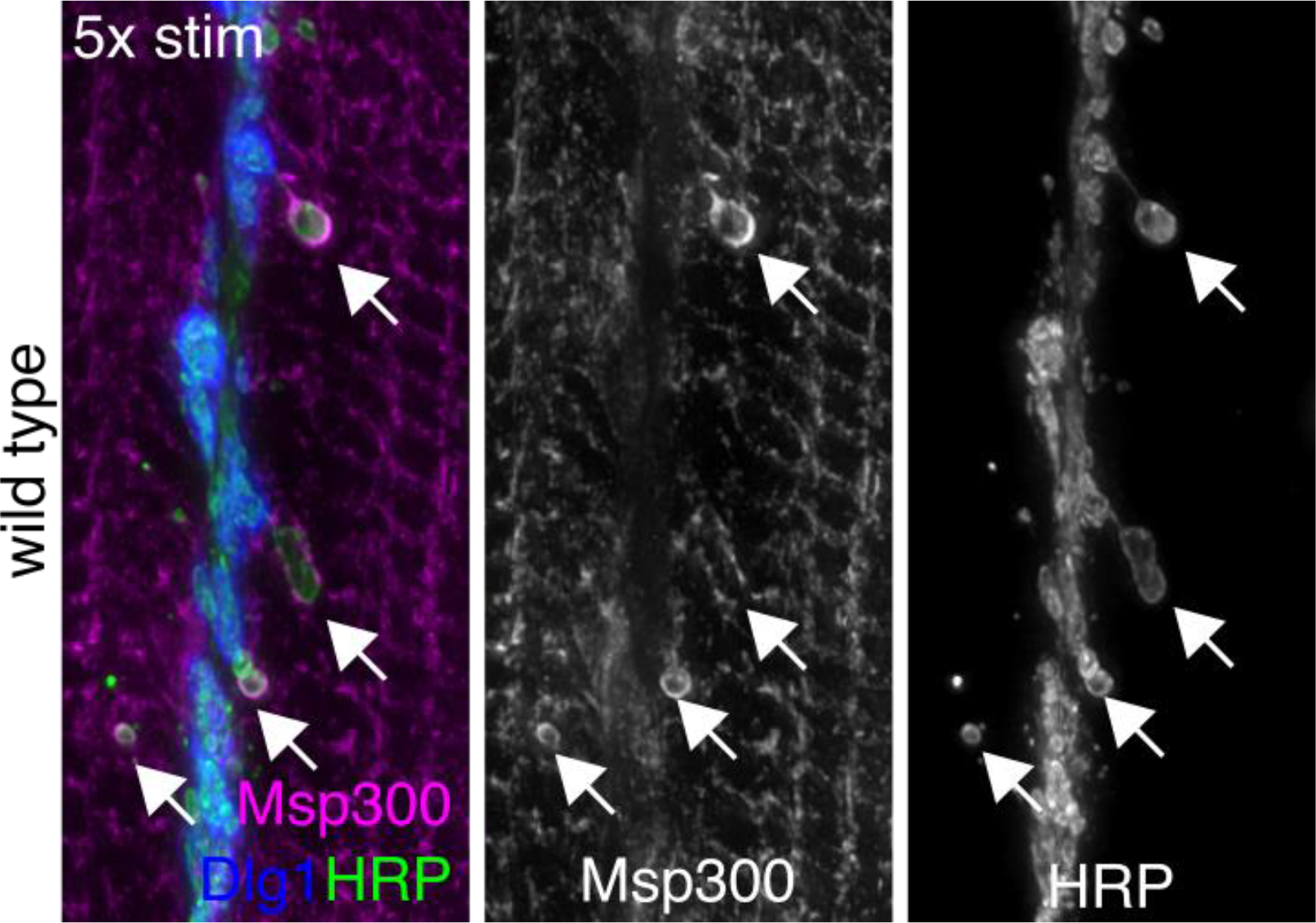
Msp300 protein is enriched around activity-induced ghost boutons. Msp300 accumulates specifically around newly formed synapses (ghost boutons). Maximum z-projection of spinning disk confocal images show strong Msp300 immunofluorescence around ghost boutons (arrows) after spaced KCl stimulation, but not around mature synapses (labeled by Dlg1, blue).

**Figure S2.**
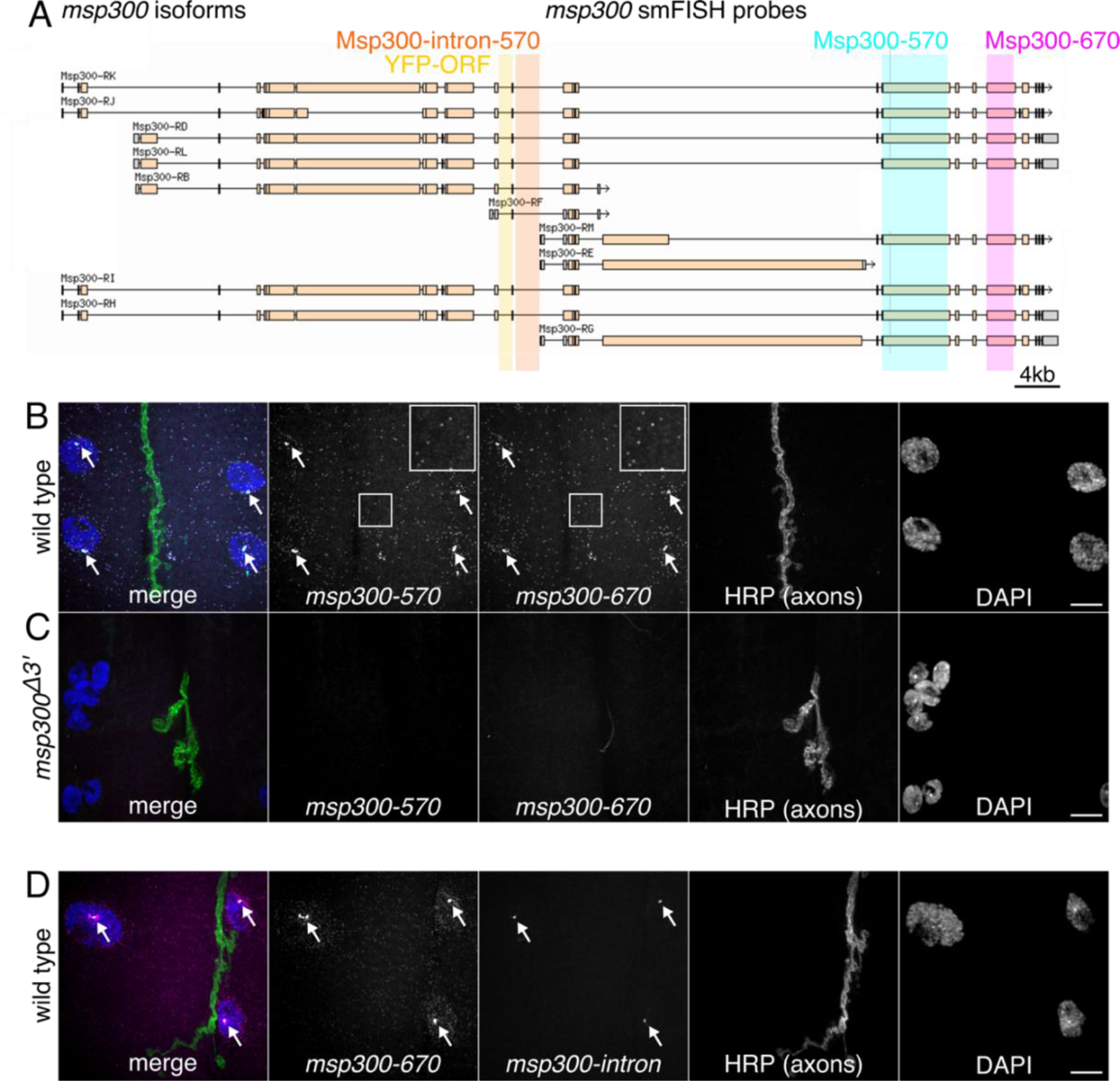
smFISH probes for *msp300* have high detection efficiency and specificity. (A) Schematic of *msp300* mRNA isoforms showing the position of smFISH probes and the YFP protein trap insertion used in this study. smFISH probes were designed to target large exons at the 3’ end of the transcript that are common to most isoforms (turquoise and magenta boxes), with dyes that have easily distinguishable fluorescence emission spectra, i.e., Quasar-570 (Msp300-570) and Quasar-670 (Msp300-670). (B) A two-color labelling experiment shows that signal from both *msp300* smFISH exon probes appear as bright punctae throughout the muscle cytoplasm with brighter transcription foci (arrows) in the larval NMJ (top row). The cytosolic spots have a uniform intensity and the majority of spots are detected in both channels (insets), indications that the signal arises from single molecules and that the detection efficiency is high. (C) smFISH probes for *msp300* don’t show any off-target binding. Punctate smFISH signal is not observed in an Msp300 mutant (*msp300∆3’*) that was hybridized, and imaged under identical acquisition settings as in B. (D) Intron/exon smFISH experiment shows that the large nuclear foci in (B) are primary *msp300* transcripts. Signal from an smFISH probe targeting intronic sequence overlaps with the Msp300-670 exon probe, but does not label mature mRNA in the nucleus or cytoplasm. Images are maximum intensity projections of spinning disk confocal sections; scale bars = 10μm.

**Figure S3.**
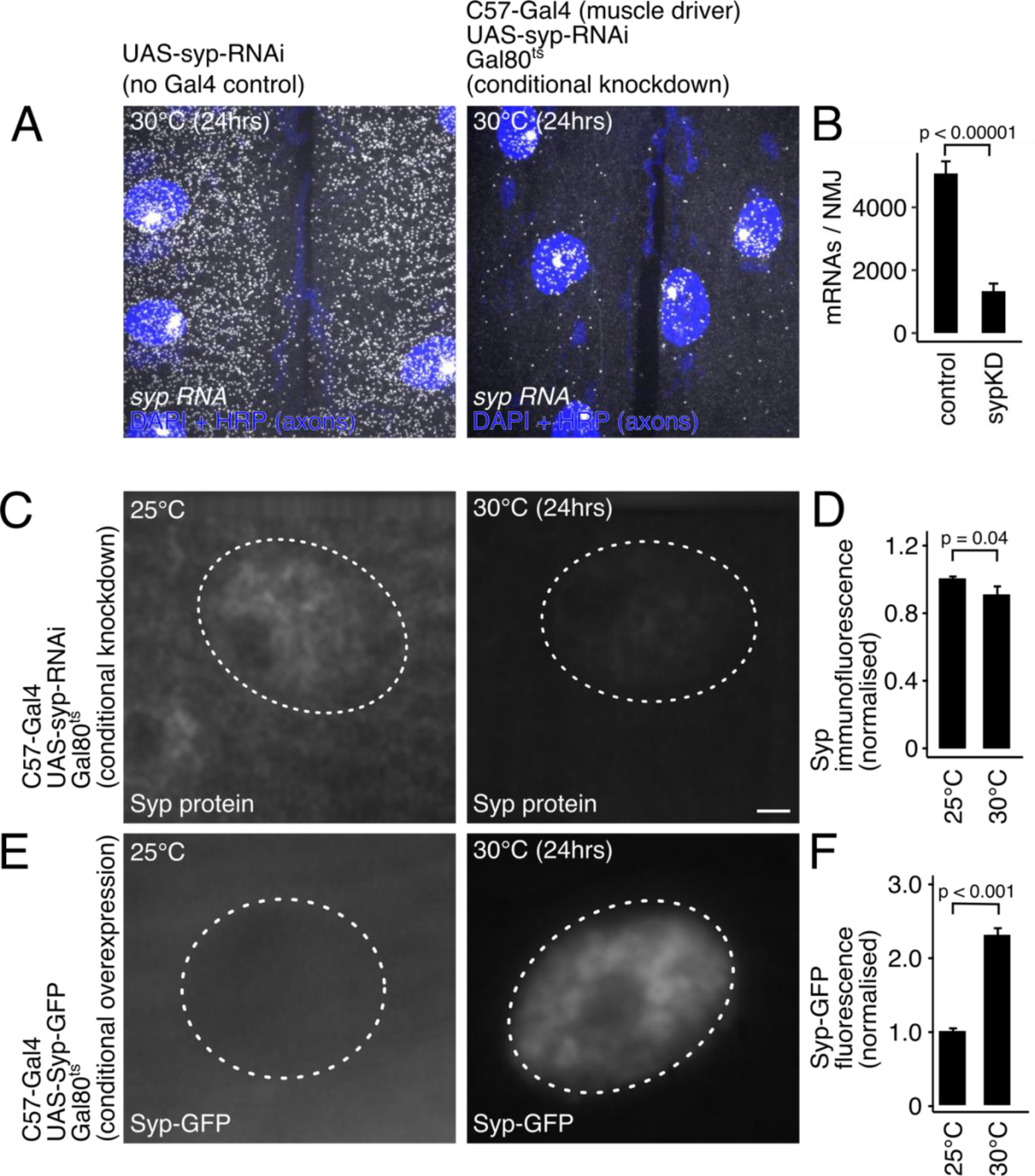
Quantification of conditional *syp* knockdown and overexpression. (A) *syp* mRNA levels are dramatically decreased by conditional knockdown, i.e., transferring 3rd instar larvae (tub-Gal80^ts^; C57-Gal4>Syp-RNAi) to the restrictive temperature (30ºC) for 24hrs. Max z-projections of spinning disk confocal images showing *syp* mRNA detection with smFISH. (B) Quantification of smFISH images from control and *syp* knockdown NMJs show a significant decrease in *syp* mRNA levels (mean ± SEM; student’s t-test; N=5 NMJs/condition). (C) Immunofluorescence images show that Syp protein levels are conditionally reduced by expressing *syp* RNAi specifically during 3^rd^ larval instar stage. (D) Quantification of average Syp protein levels shows a significant reduction in Syp protein grown at the Gal 80 restrictive temperature (mean ± SEM; student’s t-test; N=10 NMJs/condition, average of 3 nuclei/NMJ). (E) Immunofluorescence images showing that Syp-GFP is conditionally overexpressed in muscle by shifting larvae to the restrictive temperature. (F) Quantification of Syp-GFP fluorescence shows that protein levels are significantly elevated at the Gal80 restrictive temperature (mean ± SEM; student’s t-test; N=10 NMJs/condition, average of 3 nuclei/NMJ; scale bar = 2μm).

**Figure S4.**
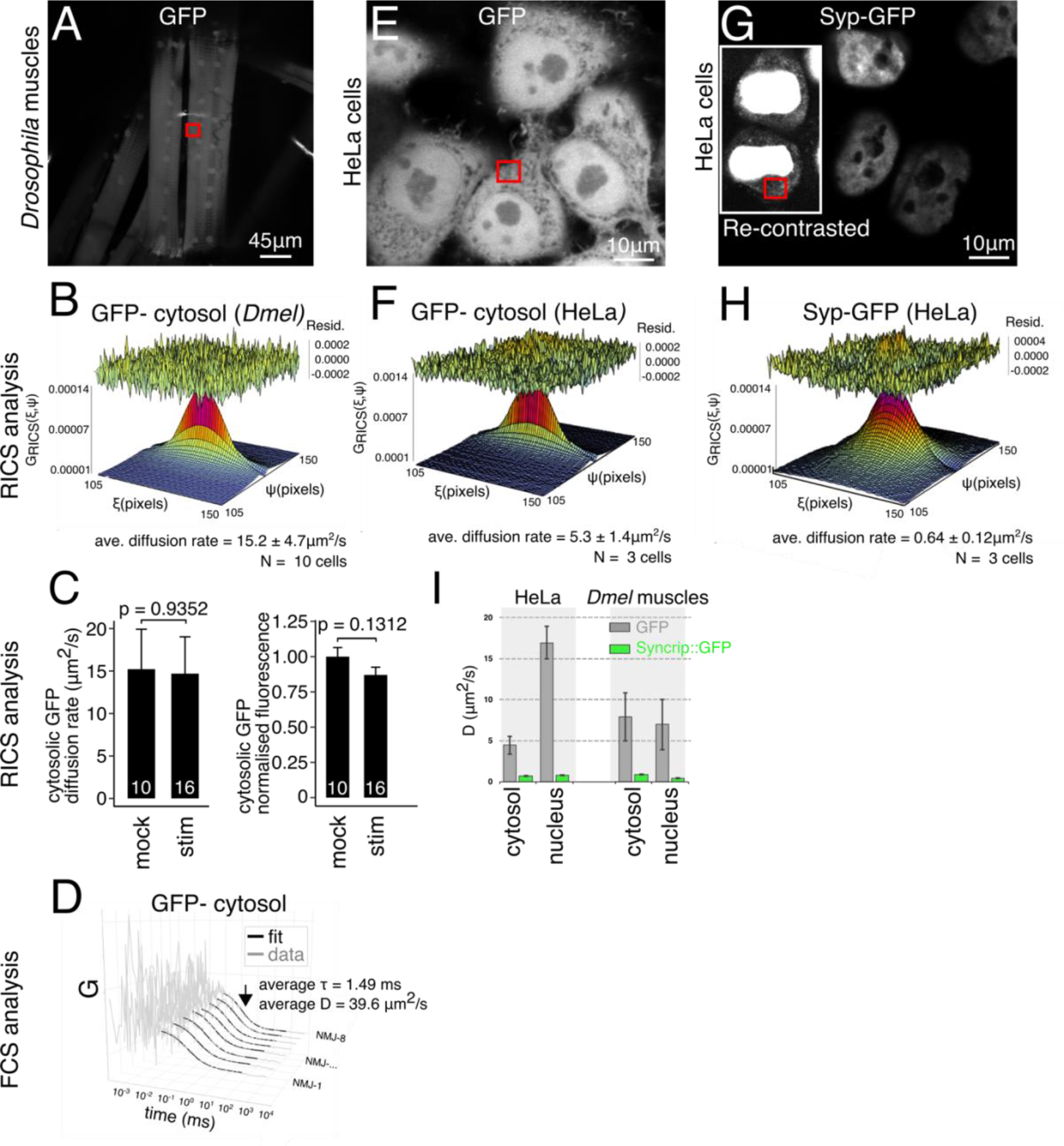
Syp-GFP diffusion is over 10 times slower than GFP in *Drosophila muscle* and human cells. (A) GFP expression in live larval muscle cells, max z-projection of confocal image. Red box indicates the region of interest (ROI) imaged for RICS data acquisition. (B) Representative fit of the autocorrelation function from the ROI in (A), used to estimate the rate of diffusion. (C) Cytosolic GFP diffusion and fluorescence intensity in larval muscles are unaffected by KCl stimulation. (D) Representative fluorescence correlation spectroscopy (FCS) curves acquired from the ROI in A as an independent measure of diffusion rate. The curves were fit using a standard 3D, 1-species diffusion model and showed diffusion rates similar to RICS measurements. (E) Live, confocal image of doxycycline-induced GFP expression in HeLa cells. (F) Representative fit of the autocorrelation function from the ROI labeled in panel (E). (G) Live, confocal image of doxycycline-induced human Syp-GFP expression in HeLa cells. Inset shows the same image re-contrasted to reveal cytosolic Syp expression, which is much lower than nuclear expression. (H) Representative fit of the autocorrelation function from the ROI labeled in (G). (I) Comparison of average diffusion rates for GFP and Syp-GFP in different compartments of *Drosophila* muscle cells and HeLa cells types.

**Figure S5.**
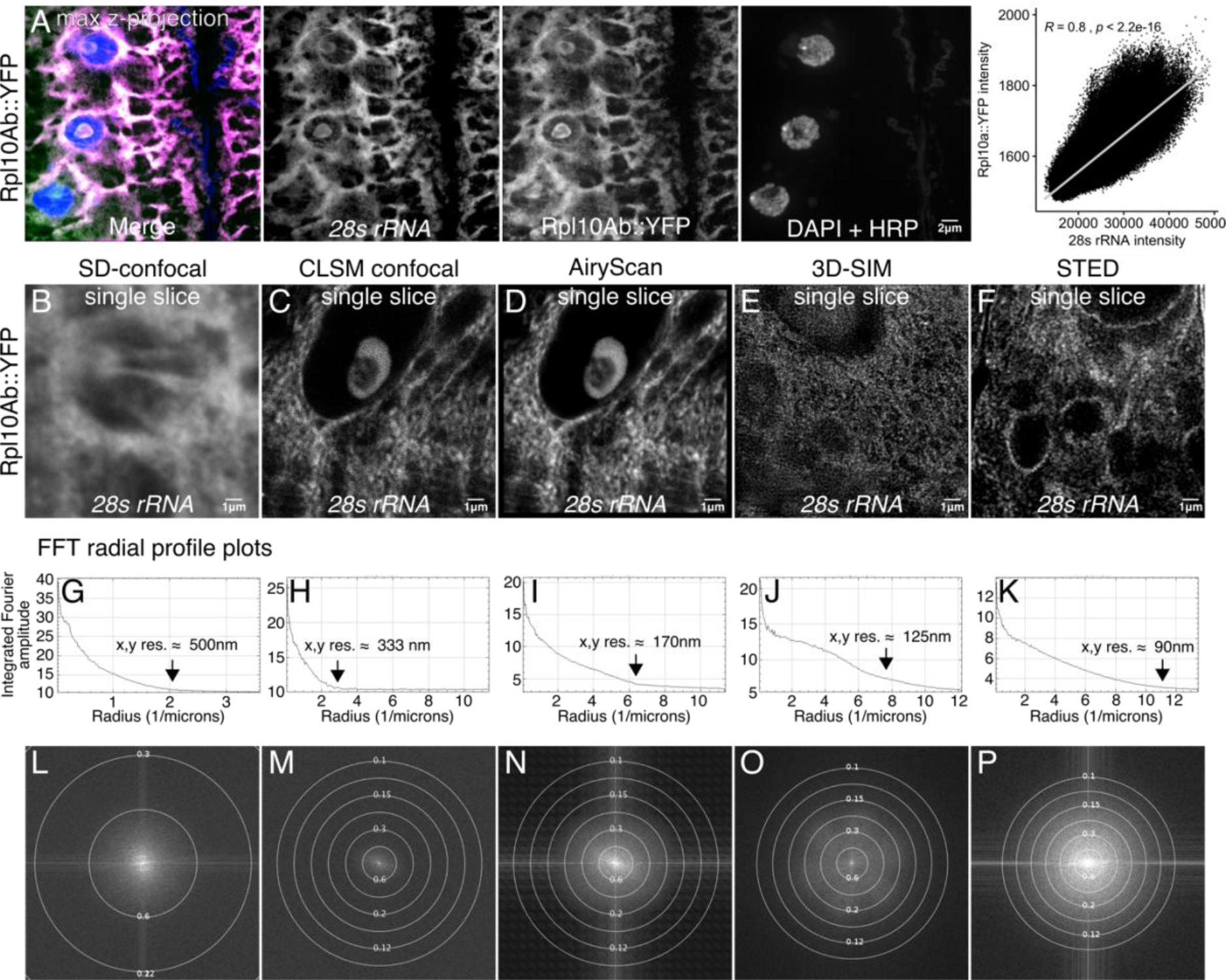
Detection of ribosome clusters using an smFISH probe targeting 28s rRNA. (A) Signal from the 28s rRNA smFISH probe overlaps almost completely with signal from Rpl10Ab::YFP protein trap in larval muscles. Images are max z-projections from a spinning disk confocal. Quantification of individual pixel intensity for 28s rRNA and Rpl10Ab::YFP signal shows highly significant correlation (Pearson correlation). (B-F) AiryScan confocal provides adequate resolution of ribosome clusters in the larval NMJ. Comparison of different confocal and super resolution microscopy techniques for imaging ribosome clusters in the NMJ with the 28s rRNA smFISH probe. The CLSM confocal image (C) is from a single AiryScan detector of the same image that has been processed in (D). The large pinhole (~3.5AU) makes for an image with suboptimal resolution relative to a properly acquired LSM image. (E) It is not possible to achieve the highest resolution enhancement with 3D-SIM due to high background signal that interferes with stripe contrast from the projected SIM pattern. (F) With STED, we were able to achieve lateral resolution below 100nm, revealing small clusters of ribosomes within the nuclear envelope and at the post synaptic density. Of the techniques tested STED provided the best resolution, however excessive photon damage from the STED laser prohibited 3D sectioning. Therefore, we chose to perform co-localisation experiments using the AiryScan system, which still enabled visualization of ribosome clusters that were observed in STED. (G-P) FFT radial profile plots provide a quantitative estimate of the resolution of each system.

## References

Ataman, B., J. Ashley, M. Gorczyca, P. Ramachandran, W. Fouquet, S.J. Sigrist, and V. Budnik. 2008. Rapid activity-dependent modifications in synaptic structure and function require bidirectional Wnt signaling. Neuron. 57:705–718.

Ball, G., J. Demmerle, R. Kaufmann, I. Davis, I.M. Dobbie, and L. Schermelleh. 2015. SIMcheck: a Toolbox for Successful Super-resolution Structured Illumination Microscopy. Scientific reports. 5:15915.

Bannai, H., K. Fukatsu, A. Mizutani, T. Natsume, S. Iemura, T. Ikegami, T. Inoue, and K. Mikoshiba. 2004. An RNA-interacting protein, SYNCRIP (heterogeneous nuclear ribonuclear protein Q1/NSAP1) is a component of mRNA granule transported with inositol 1,4,5-trisphosphate receptor type 1 mRNA in neuronal dendrites. The Journal of biological chemistry. 279:53427–53434.

Blunk, A.D., Y. Akbergenova, R.W. Cho, J. Lee, U. Walldorf, K. Xu, G. Zhong, X. Zhuang, and J.T. Littleton. 2014. Postsynaptic actin regulates active zone spacing and glutamate receptor apposition at the Drosophila neuromuscular junction. Molecular and cellular neurosciences. 61:241–254.

Brown, C.M., R.B. Dalal, B. Hebert, M.A. Digman, A.R. Horwitz, and E. Gratton. 2008. Raster image correlation spectroscopy (RICS) for measuring fast protein dynamics and concentrations with a commercial laser scanning confocal microscope. Journal of microscopy. 229:78–91.

Buxbaum, A.R., B. Wu, and R.H. Singer. 2014. Single beta-actin mRNA detection in neurons reveals a mechanism for regulating its translatability. Science. 343:419–422.

Chen, H.H., H.I. Yu, W.C. Chiang, Y.D. Lin, B.C. Shia, and W.Y. Tarn. 2012. hnRNP Q regulates Cdc42-mediated neuronal morphogenesis. Molecular and cellular biology. 32:2224–2238.

Chen, X., R. Rahman, F. Guo, and M. Rosbash. 2016. Genome-wide identification of neuronal activity-regulated genes in Drosophila. eLife. 5:e19942.

Cottrell, J.R., E. Borok, T.L. Horvath, and E. Nedivi. 2004. CPG2: a brain- and synapse-specific protein that regulates the endocytosis of glutamate receptors. Neuron. 44:677–690.

Dieterich, D.C., and M.R. Kreutz. 2016. Proteomics of the Synapse – A Quantitative Approach to Neuronal Plasticity. Molecular & Cellular Proteomics. 15:368–381.

Digman, M.A., P.W. Wiseman, A.R. Horwitz, and E. Gratton. 2009. Detecting protein complexes in living cells from laser scanning confocal image sequences by the cross correlation raster image spectroscopy method. Biophysical journal. 96:707–716.

Duning, K., F. Buck, A. Barnekow, and J. Kremerskothen. 2008. SYNCRIP, a component of dendritically localized mRNPs, binds to the translation regulator BC200 RNA. Journal of neurochemistry. 105:351–359.

Dupre, N., F. Gros-Louis, N. Chrestian, S. Verreault, D. Brunet, D. de Verteuil, B. Brais, J.P. Bouchard, and G.A. Rouleau. 2007. Clinical and genetic study of autosomal recessive cerebellar ataxia type 1. Ann Neurol. 62:93–98.

Eom, T., L.N. Antar, R.H. Singer, and G.J. Bassell. 2003. Localization of a beta-actin messenger ribonucleoprotein complex with zipcode-binding protein modulates the density of dendritic filopodia and filopodial synapses. The Journal of neuroscience : the official journal of the Society for Neuroscience. 23:10433–10444.

Evangelista, M., B.M. Klebl, A.H. Tong, B.A. Webb, T. Leeuw, E. Leberer, M. Whiteway, D.Y. Thomas, and C. Boone. 2000. A role for myosin-I in actin assembly through interactions with Vrp1p, Bee1p, and the Arp2/3 complex. The Journal of cell biology. 148:353–362.

Gama, M.T., G. Houle, A. Noreau, A. Dionne-Laporte, P.A. Dion, G.A. Rouleau, O.G. Barsottini, and J.L. Pedroso. 2016. SYNE1 mutations cause autosomal-recessive ataxia with retained reflexes in Brazilian patients. Movement disorders : official journal of the Movement Disorder Society. 31:1754–1756.

Gama, M.T.D., C.C. Piccinin, T.J.R. Rezende, P.A. Dion, G.A. Rouleau, M.C. Franca Junior, O.G.P. Barsottini, and J.L. Pedroso. 2018. Multimodal neuroimaging analysis in patients with SYNE1 Ataxia. J Neurol Sci. 390:227–230.

Gros-Louis, F., N. Dupre, P. Dion, M.A. Fox, S. Laurent, S. Verreault, J.R. Sanes, J.P. Bouchard, and G.A. Rouleau. 2007. Mutations in SYNE1 lead to a newly discovered form of autosomal recessive cerebellar ataxia. Nature genetics. 39:80–85.

Gura Sadovsky, R., S. Brielle, D. Kaganovich, and J.L. England. 2017. Measurement of Rapid Protein Diffusion in the Cytoplasm by Photo-Converted Intensity Profile Expansion. Cell reports. 18:2795–2806.

Guzowski, J.F., G.L. Lyford, G.D. Stevenson, F.P. Houston, J.L. McGaugh, P.F. Worley, and C.A. Barnes. 2000. Inhibition of activity-dependent arc protein expression in the rat hippocampus impairs the maintenance of long-term potentiation and the consolidation of long-term memory. The Journal of neuroscience : the official journal of the Society for Neuroscience. 20:3993–4001.

Halstead, J.M., Y.Q. Lin, L. Durraine, R.S. Hamilton, G. Ball, G.G. Neely, H.J. Bellen, and I. Davis. 2014. Syncrip/hnRNP Q influences synaptic transmission and regulates BMP signaling at the Drosophila neuromuscular synapse. Biology open. 3:839–849.

Harris, K.P., and J.T. Littleton. 2015. Transmission, Development, and Plasticity of Synapses. Genetics. 201:345–375.

Hayashi, R., S.M. Wainwright, S.J. Liddell, S.M. Pinchin, S. Horswell, and D. Ish-Horowicz. 2014. A genetic screen based on in vivo RNA imaging reveals centrosome-independent mechanisms for localizing gurken transcripts in Drosophila. G3 (Bethesda). 4:749–760.

Jaramillo, A.M., T.T. Weil, J. Goodhouse, E.R. Gavis, and T. Schupbach. 2008. The dynamics of fluorescently labeled endogenous gurken mRNA in Drosophila. Journal of cell science. 121:887–894.

Katz, Z.B., B.P. English, T. Lionnet, Y.J. Yoon, N. Monnier, B. Ovryn, M. Bathe, and R.H. Singer. 2016. Mapping translation ‘hot-spots’ in live cells by tracking single molecules of mRNA and ribosomes. eLife. 5.

Kim, D.Y., W. Kim, K.H. Lee, S.H. Kim, H.R. Lee, H.J. Kim, Y. Jung, J.H. Choi, and K.T. Kim. 2013. hnRNP Q regulates translation of p53 in normal and stress conditions. Cell Death Differ. 20:226–234.

Kim, J.H., K.Y. Paek, S.H. Ha, S. Cho, K. Choi, C.S. Kim, S.H. Ryu, and S.K. Jang. 2004. A cellular RNA-binding protein enhances internal ribosomal entry site-dependent translation through an interaction downstream of the hepatitis C virus polyprotein initiation codon. Molecular and cellular biology. 24:7878–7890.

Korobchevskaya, K., C.B. Lagerholm, H. Colin-York, and M. Fritzsche. 2017. Exploring the Potential of Airyscan Microscopy for Live Cell Imaging. Photonics. 4.

Kuchler, L., A.K. Giegerich, L.K. Sha, T. Knape, M.S. Wong, K. Schroder, R.P. Brandes, H. Heide, I. Wittig, B. Brune, and A. von Knethen. 2014. SYNCRIP-dependent Nox2 mRNA destabilization impairs ROS formation in M2-polarized macrophages. Antioxid Redox Signal. 21:2483–2497.

Lee, K.H., K.C. Woo, D.Y. Kim, T.D. Kim, J. Shin, S.M. Park, S.K. Jang, and K.T. Kim. 2012. Rhythmic interaction between Period1 mRNA and hnRNP Q leads to circadian time-dependent translation. Molecular and cellular biology. 32:717–728.

Lelieveld, S.H., M.R. Reijnders, R. Pfundt, H.G. Yntema, E.J. Kamsteeg, P. de Vries, B.B. de Vries, M.H. Willemsen, T. Kleefstra, K. Lohner, M. Vreeburg, S.J. Stevens, I. van der Burgt, E.M. Bongers, A.P. Stegmann, P. Rump, T. Rinne, M.R. Nelen, J.A. Veltman, L.E. Vissers, H.G. Brunner, and C. Gilissen. 2016. Meta-analysis of 2,104 trios provides support for 10 new genes for intellectual disability. Nature neuroscience. 19:1194–1196.

Lowe, N., J.S. Rees, J. Roote, E. Ryder, I.M. Armean, G. Johnson, E. Drummond, H. Spriggs, J. Drummond, J.P. Magbanua, H. Naylor, B. Sanson, R. Bastock, S. Huelsmann, V. Trovisco, M. Landgraf, S. Knowles-Barley, J.D. Armstrong, H. White-Cooper, C. Hansen, R.G. Phillips, U.K.D.P.T.S. Consortium, K.S. Lilley, S. Russell, and D. St Johnston. 2014. Analysis of the expression patterns, subcellular localisations and interaction partners of Drosophila proteins using a pigP protein trap library. Development. 141:3994–4005.

Madabhushi, R., and T.-K. Kim. 2018. Emerging themes in neuronal activity-dependent gene expression. Molecular and Cellular Neuroscience. 87:27–34.

Mademan, I., F. Harmuth, I. Giordano, D. Timmann, S. Magri, T. Deconinck, J. Claassen, D. Jokisch, G. Genc, D. Di Bella, S. Romito, R. Schule, S. Zuchner, F. Taroni, T. Klockgether, L. Schols, P. De Jonghe, P. Bauer, E. Consortium, J. Baets, and M. Synofzik. 2016. Multisystemic SYNE1 ataxia: confirming the high frequency and extending the mutational and phenotypic spectrum. Brain. 139:e46.

Matsuda, A., L. Schermelleh, Y. Hirano, T. Haraguchi, and Y. Hiraoka. 2018. Accurate and fiducial-marker-free correction for three-dimensional chromatic shift in biological fluorescence microscopy. Scientific reports. 8:7583.

McAllister, A.K. 2007. Dynamic Aspects of CNS Synapse Formation. Annual review of neuroscience. 30:425–450.

McDermott, S.M., L. Yang, J.M. Halstead, R.S. Hamilton, C. Meignin, and I. Davis. 2014. Drosophila Syncrip modulates the expression of mRNAs encoding key synaptic proteins required for morphology at the neuromuscular junction. Rna. 20:1593–1606.

Menon, K.P., R.A. Carrillo, and K. Zinn. 2013. Development and plasticity of the Drosophila larval neuromuscular junction. Wiley interdisciplinary reviews. Developmental biology. 2:647–670.

Morel, V., S. Lepicard, A.N. Rey, M.L. Parmentier, and L. Schaeffer. 2014. Drosophila Nesprin-1 controls glutamate receptor density at neuromuscular junctions. Cellular and molecular life sciences : CMLS. 71:3363–3379.

Munno David, W., and I. Syed Naweed. 2004. Synaptogenesis in the CNS: An Odyssey from Wiring Together to Firing Together. The Journal of physiology. 552:1–11.

Noreau, A., C.V. Bourassa, A. Szuto, A. Levert, S. Dobrzeniecka, J. Gauthier, S. Forlani, A. Durr, M. Anheim, G. Stevanin, A. Brice, J.P. Bouchard, P.A. Dion, N. Dupre, and G.A. Rouleau. 2013. SYNE1 mutations in autosomal recessive cerebellar ataxia. JAMA Neurol. 70:1296–1231.

Packard, M., V. Jokhi, B. Ding, C. Ruiz-Canada, J. Ashley, and V. Budnik. 2015. Nucleus to Synapse Nesprin1 Railroad Tracks Direct Synapse Maturation through RNA Localization. Neuron. 86:1015–1028.

Rossow, M.J., J.M. Sasaki, M.A. Digman, and E. Gratton. 2010. Raster image correlation spectroscopy in live cells. Nature protocols. 5:1761–1774.

Rubin, G.M., L. Hong, P. Brokstein, M. Evans-Holm, E. Frise, M. Stapleton, and D.A. Harvey. 2000. A Drosophila complementary DNA resource. Science. 287:2222–2224.

Santangelo, L., G. Giurato, C. Cicchini, C. Montaldo, C. Mancone, R. Tarallo, C. Battistelli, T. Alonzi, A. Weisz, and M. Tripodi. 2016. The RNA-Binding Protein SYNCRIP Is a Component of the Hepatocyte Exosomal Machinery Controlling MicroRNA Sorting. Cell reports. 17:799–808.

Scofield, S.R., and W.Y. Chooi. 1982. Structure of ribosomes and ribosomal subunits of Drosophila. Mol Gen Genet. 187:37–41.

Spence, E.F., and S.H. Soderling. 2015. Actin Out: Regulation of the Synaptic Cytoskeleton. The Journal of biological chemistry. 290:28613–28622.

Steward, O., C.S. Wallace, G.L. Lyford, and P.F. Worley. 1998. Synaptic activation causes the mRNA for the IEG Arc to localize selectively near activated postsynaptic sites on dendrites. Neuron. 21:741–751.

Steward, O., and P.F. Worley. 2001. A cellular mechanism for targeting newly synthesized mRNAs to synaptic sites on dendrites. Proceedings of the National Academy of Sciences of the United States of America. 98:7062–7068.

Stroud, M.J., W. Feng, J. Zhang, J. Veevers, X. Fang, L. Gerace, and J. Chen. 2017. Nesprin 1alpha2 is essential for mouse postnatal viability and nuclear positioning in skeletal muscle. The Journal of cell biology. 216:1915–1924.

Suster, M.L., L. Seugnet, M. Bate, and M.B. Sokolowski. 2004. Refining GAL4-driven transgene expression in Drosophila with a GAL80 enhancer-trap. Genesis. 39:240–245.

Synofzik, M., K. Smets, M. Mallaret, D. Di Bella, C. Gallenmuller, J. Baets, M. Schulze, S. Magri, E. Sarto, M. Mustafa, T. Deconinck, T. Haack, S. Zuchner, M. Gonzalez, D. Timmann, C. Stendel, T. Klopstock, A. Durr, C. Tranchant, M. Sturm, W. Hamza, L. Nanetti, C. Mariotti, M. Koenig, L. Schols, R. Schule, P. de Jonghe, M. Anheim, F. Taroni, and P. Bauer. 2016. SYNE1 ataxia is a common recessive ataxia with major non-cerebellar features: a large multi-centre study. Brain. 139:1378–1393.

Titlow, J.S., and R.L. Cooper. 2018. Glutamatergic Synthesis, Recycling, and Receptor Pharmacology at Drosophila and Crustacean Neuromuscular Junctions. In Biochemical Approaches for Glutamatergic Neurotransmission. S. Parrot and L. Denoroy, editors. Springer New York, New York, NY. 263–291.

Titlow, J.S., L. Yang, R.M. Parton, A. Palanca, and I. Davis. 2018. Super-Resolution Single Molecule FISH at the Drosophila Neuromuscular Junction. Methods in molecular biology. 1649:163–175.

Tratnjek, L., M. Zivin, and G. Glavan. 2017. Synaptotagmin 7 and SYNCRIP proteins are ubiquitously expressed in the rat brain and co-localize in Purkinje neurons. J Chem Neuroanat. 79:12–21.

Ukken, F.P., J.J. Bruckner, K.L. Weir, S.J. Hope, S.L. Sison, R.M. Birschbach, L. Hicks, K.L. Taylor, E.W. Dent, G.B. Gonsalvez, and K.M. Connor-Giles. 2016. BAR-SH3 sorting nexins are conserved interacting proteins of Nervous wreck that organize synapses and promote neurotransmission. Journal of cell science. 129:166.

Verschoor, A., S. Srivastava, R. Grassucci, and J. Frank. 1996. Native 3D structure of eukaryotic 80s ribosome: morphological homology with E. coli 70S ribosome. The Journal of cell biology. 133:495–505.

Verschoor, A., J.R. Warner, S. Srivastava, R.A. Grassucci, and J. Frank. 1998. Three-dimensional structure of the yeast ribosome. Nucleic acids research. 26:655–661.

Vincendeau, M., D. Nagel, J.K. Brenke, R. Brack-Werner, and K. Hadian. 2013. Heterogenous nuclear ribonucleoprotein Q increases protein expression from HIV-1 Rev-dependent transcripts. Virol J. 10:151.

Volk, T. 2013. Positioning nuclei within the cytoplasm of striated muscle fiber. Nucleus. 4:18–22.

Vu, L.P., C. Prieto, E.M. Amin, S. Chhangawala, A. Krivtsov, M.N. Calvo-Vidal, T. Chou, A. Chow, G. Minuesa, S.M. Park, T.S. Barlowe, J. Taggart, P. Tivnan, R.P. Deering, L.P. Chu, J.A. Kwon, C. Meydan, J. Perales-Paton, A. Arshi, M. Gonen, C. Famulare, M. Patel, E. Paietta, M.S. Tallman, Y. Lu, J. Glass, F.E. Garret-Bakelman, A. Melnick, R. Levine, F. Al-Shahrour, M. Jaras, N. Hacohen, A. Hwang, R. Garippa, C.J. Lengner, S.A. Armstrong, L. Cerchietti, G.S. Cowley, D. Root, J. Doench, C. Leslie, B.L. Ebert, and M.G. Kharas. 2017. Functional screen of MSI2 interactors identifies an essential role for SYNCRIP in myeloid leukemia stem cells. Nature genetics. 49:866–875.

Waithe, D., M.P. Clausen, E. Sezgin, and C. Eggeling. 2016. FoCuS-point: software for STED fluorescence correlation and time-gated single photon counting. Bioinformatics. 32:958–960.

Wang, S., A. Reuveny, and T. Volk. 2015. Nesprin provides elastic properties to muscle nuclei by cooperating with spectraplakin and EB1. The Journal of cell biology. 209:529–538.

West, A.E., and M.E. Greenberg. 2011. Neuronal activity-regulated gene transcription in synapse development and cognitive function. Cold Spring Harbor perspectives in biology. 3.

Wiethoff, S., J. Hersheson, C. Bettencourt, N.W. Wood, and H. Houlden. 2016. Heterogeneity in clinical features and disease severity in ataxia-associated SYNE1 mutations. J Neurol. 263:1503–1510.

Will, T.J., G. Tushev, L. Kochen, B. Nassim-Assir, I.J. Cajigas, S. Tom Dieck, and E.M. Schuman. 2013. Deep sequencing and high-resolution imaging reveal compartment-specific localization of Bdnf mRNA in hippocampal neurons. Science signaling. 6:rs16.

Zhan, H., J. Bruckner, Z. Zhang, and K. O’Connor-Giles. 2016. Three-dimensional imaging of Drosophila motor synapses reveals ultrastructural organizational patterns. Journal of neurogenetics. 30:237–246.

Zhang, X., K. Lei, X. Yuan, X. Wu, Y. Zhuang, T. Xu, R. Xu, and M. Han. 2009. SUN1/2 and Syne/Nesprin-1/2 complexes connect centrosome to the nucleus during neurogenesis and neuronal migration in mice. Neuron. 64:173–187.

Zhou, C., L. Rao, C.M. Shanahan, and Q. Zhang. 2018a. Nesprin-1/2: roles in nuclear envelope organisation, myogenesis and muscle disease. Biochemical Society transactions. 46:311.

Zhou, C., L. Rao, D.T. Warren, C.M. Shanahan, and Q. Zhang. 2018b. Mouse models of nesprin-related diseases. Biochemical Society transactions. 46:669–681.

